# High-Dimensional Sensitivity Analysis for Genomic Studies: An Adversarial Framework for Learning Worst-Case Latent Confounders

**DOI:** 10.64898/2026.05.27.728283

**Authors:** Yifan Lin, Kevin Z. Lin

## Abstract

High-dimensional genomics studies are frequently confounded by unmeasured biological processes that obscure disease-specific signals. While existing workflows can estimate these latent confounders, they fail to quantify how robust a discovery is to varying levels of hypothetical confounding. We introduce sensGAN, a deep-learning adversarial framework that systematically explores the confounding spectrum by learning “worst-case” latent variables that nullify the most gene associations under novel predictive-gain constraints. By identifying the minimum confounding strength required to explain away an observed effect, our method shifts the paradigm toward a formal, quantitative sensitivity analysis. In diverse simulations, sensGAN accurately recovers latent structures and outperforms existing methods in identifying confounder-sensitive genes. Applied to human Alzheimer’s disease microglia, our framework prioritizes robust disease pathways while successfully isolating signals driven by unmeasured co-occurring neurodegenerative pathologies. Our method is publicly available, deposited at the GitHub repository `yifanlinz/AD_sensitivity_ICML`.

## 1. Introduction

A fundamental challenge in modern genomics is the presence of unmeasured confounding variables that can obscure true biological signals. In high-dimensional single-cell studies, transcriptional changes attributed to a primary disease state may instead reflect unmeasured comorbid processes, environmental exposures, or systemic pathologies that are incompletely recorded. While conventional differential expression analyses prioritize genes based on statistical significance (Squair et al., 2021), they often fail to quantify the sensitivity of these findings to latent confounders: the more sensitive the findings are to latent confounders, the less robust the discoveries are. Computational frameworks, such as surrogate variable analysis (Leek & Storey, 2007; 2008), provide a single point estimate of latent factors to adjust for residual variation but do not explore the spectrum of hypothetical confounding. This leaves researchers without a metric to determine how “strong” a latent factor would need to be to nullify a discovery.

We argue that high-dimensional genomics requires a shift toward formal sensitivity analysis, mirroring classic statistical approaches like Cornfield’s analysis of smoking and lung cancer (Cornfield et al., 1959). Conceptually, we ask: what level of confounding strength, relative to observed covariates, is required to explain away a gene’s association with a treatment or disease label? This is a novel machine learning problem involving the optimization of “worst-case” latent variables within high-dimensional outcome spaces. By treating confounding as a spectrum rather than a fixed correction, we can distinguish between transcripts that are robust to latent biological processes and those whose significance is likely an artifact of omitted variable bias.

To address this, we introduce **sensGAN** (Sensitivity Generative Adversarial Network), a generative adversarial network for sensitivity analysis that leverages deep learning. SensGAN does not aim to identify a unique causal confounder; rather, it characterizes the space of latent variables that are sufficiently predictive to explain away an observed association. SensGAN learns worst-case latent confounders (also called “unmeasured confounders” in other literature) by constraining their predictive gains on both the treatment and outcome, identifying the weakest confounding strength required to nullify each gene’s association with the disease.

### 1.1. Scientific relevance

The necessity of sensitivity analysis is most evident in the study of complex disorders like Alzheimer’s disease (AD). A major computational obstacle in uncovering “direct” disease signals is that donors frequently harbor co-occurring neurodegenerative diseases (co-pathologies). For instance, in late-onset AD, as few as 31% of cases are described by purely AD-specific signatures, with the remainder displaying complex comorbidity patterns that potentially confound existing single-cell analyses (Robinson et al., 2023).

To illustrate the clinicial relevance of understanding the role of co-pathologies, we perform a preliminary analysis of a cohort of 4,168 deceased donors from the National Alzheimer’s Coordinating Center (NACC), all of whom were staged for AD and 5 other co-pathologies (Fig. 1A). Our analysis reinforces a prevailing scientific hypothesis that as primary disease severity increases, donors exhibit an increasing burden of these co-pathologies, which significantly impacts clinical outcomes and donor lifespan (Spina et al., 2021). Our results show that as the severity of AD increases, an increasing percentage of those donors have more co-pathologies (Fig. 1B).

**Figure 1.**
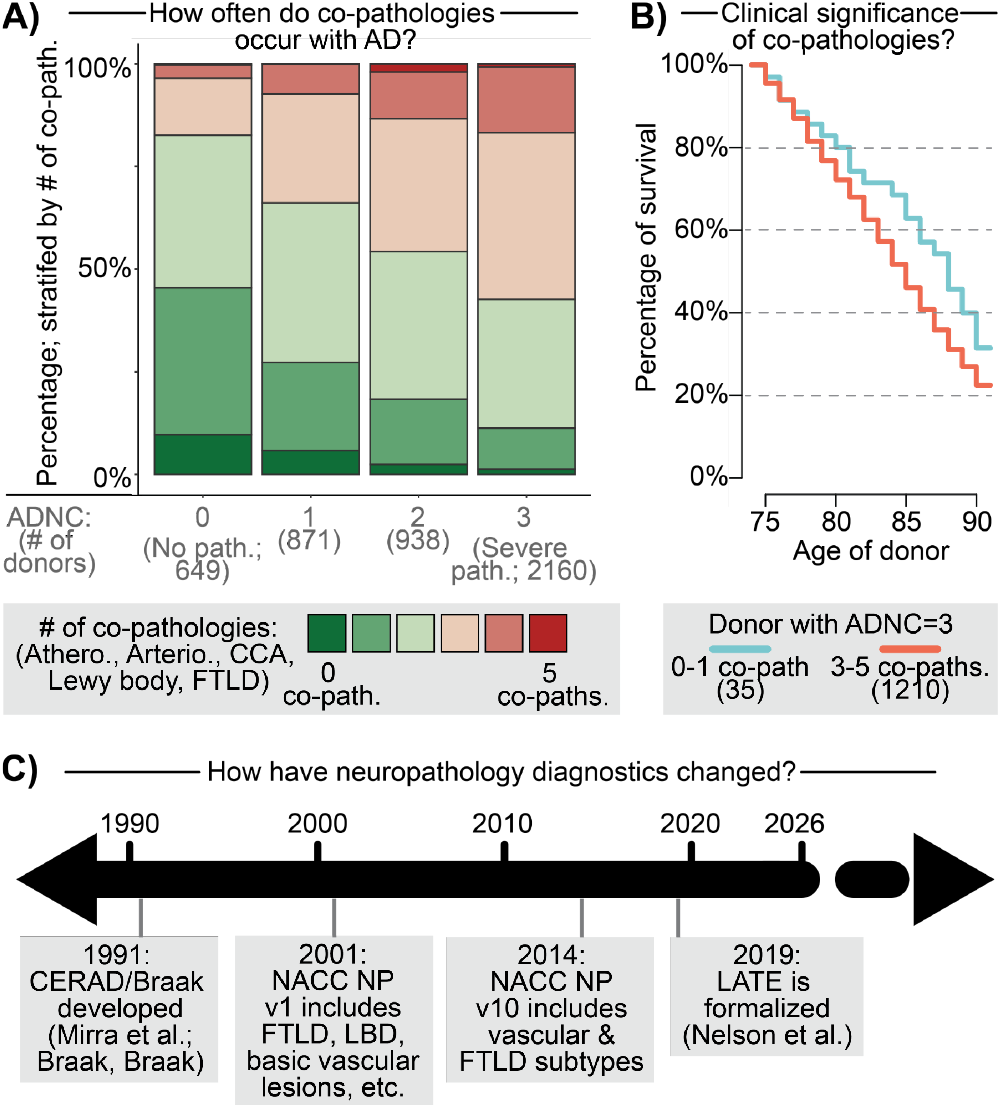
Clinical impact of latent confounders in AD research and sensGAN overview to learn them. **A)** Percentage of donors with number of co-occurring neurodegenerative disease, stratified by AD severity (ADNC score: 0 being no pathologies, 3 being severe). The number of donors in the NACC cohort in each stratum is marked. **B)** Survival curve among donors at age-at-death ≥ 75 with severe AD pathologies, stratified by whether they have 0/1 co-pathologies or 3+ co-pathologies. **C)** Timeline of evolving AD and related co-occurring neurodegenerative diseases were standardized among clinical diagnostics of post-mortem brain tissue.

**Figure 2.**
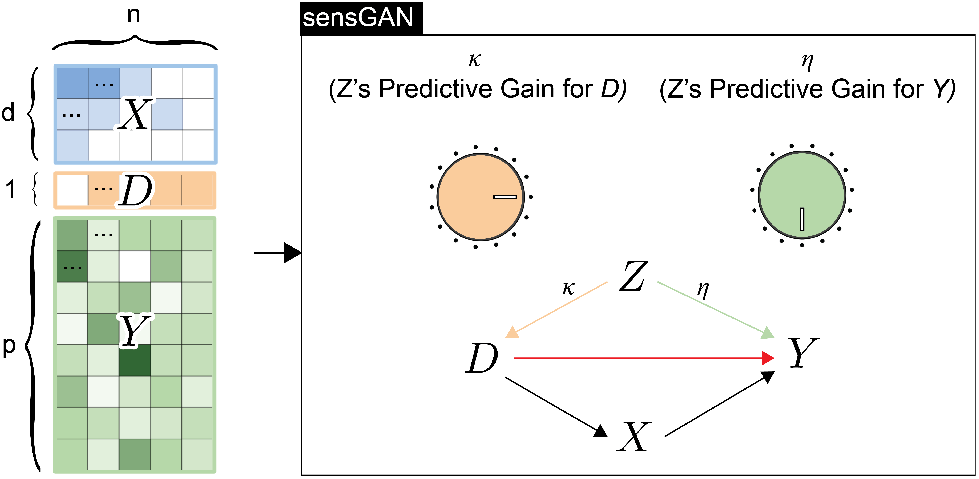
sensGAN core idea. As illustrated, unobserved confounders may simultaneously influence both the treatment *D* and the outcome *Y*, leading to spurious associations. sensGAN parameterizes this confounding strength using two normalized predictive-gain knobs and systematically explores the resulting confounding scenarios.

The challenge for computational biologists is that the clinical definitions for these co-pathologies are not static; they undergo major updates roughly every decade as scientific understanding evolves (Mirra et al., 1991; Montine et al., 2012; Nelson et al., 2019). Although many of the cited studies suggest that adjusting for co-occurring neurodegenerative disease is important to uncover the “pure” AD transcriptomic changes, the difficulty lies in the evolving definitions of these co-pathologies (Jack Jr et al., 2024). For instance, Figure 1C summarizes how the standard clinical co-pathology staging criteria evolved over the decades of AD research (Mirra et al., 1991; Braak & Braak, 1991; Montine et al., 2012; Nelson et al., 2019). Given this ever-changing landscape, how do we ensure that our single-cell findings today are robust to possible confounding co-occurring neurodegenerative disease that we might formalize in the future?

With this context, our goal for sensGAN is to provide a “future-looking” framework for differential expression. Rather than relying solely on currently formalized staging, our sensitivity analysis offers two critical advantages:

- **(Goal #1) Make robust discoveries even when important confounders are unmeasured or unknown**: SensGAN enables a sensitivity analysis with respect to latent confounders by explicitly constructing plausible worst-case latent confounders under controlled confounding strength to identify the one that nullifies the largest number of differentially expressed genes. Genes that remain significant under these worst-case scenarios are therefore robust to a broad class of latent confounding effects.
- **(Goal #2) Use the learned confounders to suggest what hidden biology might exist**: SensGAN learns explicit latent confounder representations that may reflect underlying biological processes not yet measured or formally defined. Although these latent variables are not assumed to correspond to any known pathology, their values can suggest the presence and functional impact of latent biological drivers, providing hypotheses for future experimental or clinical investigation.

### 1.2. Existing computational methods and their limitations

We model log-normalized, pseudo-bulked gene expression vector *Y* ∈ ℕ^*n×p*^ (i.e., the “outcome” across *n* donors and *p* genes), treatment status *D* ∈ {0, 1} ^*n×*1^ (i.e., the case-control status), and the *d* measured covariates *X* ∈ ℝ^*n×d*^. Following standard Generalized Linear Model (GLM) frameworks commonly used to model gene expression vector (Sarkar & Stephens, 2021), we assume:

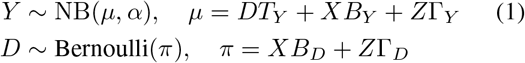

Our primary objectives are to estimate the direct effect *T*_*Y*_ ∈ ℝ^1*×p*^ of *D* on *Y* (Goal #1) and the *k*-dimensional latent factor matrix *Z* ∈ ℝ^*n×k*^ (Goal #2). Here, 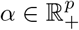 denotes gene-specific overdispersions, and *B*, Γ represent effect matrices for observed and latent variables, respectively, for either the outcome or treatment, depending on the subscript. In practice, *Y* is often derived from single-cell RNA-seq data, where we aggregate the expression profile across all cells of a particular cell type of interest from each donor, and then log-normalize the expression profile across *p* highly variable genes of interest.

However, since *Z* is unobserved, typical DE methods are forced to model the data *Y* using an “unadjusted” model,

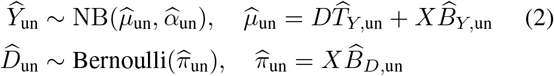

We highlight existing computational frameworks below for handling latent confounders *Z* in this setting, which motivate our proposed method.

Existing frameworks for latent confounding generally fall into two categories. Omitted Variable Bias (OVB) methods quantify the strength required for a latent factor to nullify a treatment-outcome relation (Cinelli & Hazlett, 2020; Veitch & Zaveri, 2020). However, these are largely restricted to linear models and do not scale to the non-linear, high-dimensional outcome spaces (e.g., thousands of genes modeled via Negative Binomial (NB) distributions) typical of single-cell genomics. Conversely, surrogate-variable approaches, such as causarray (Du et al., 2026) and RUVr (Risso et al., 2014), estimate latent factors to adjust for residual variation. These methods are conceptually related to PCA-based approaches, as they seek latent factors that explain as much residual variability as possible after accounting for measured covariates. Yet, these provide only point estimates rather than exploring the spectrum of hypothetical confounding. Furthermore, they often assume that latent factors are independent of the treatment, making them poorly suited to capture neurodegenerative diseases that co-occur with primary disease states like AD. See Appendix S2 for additional discussion.

## 2. Overview of sensGAN

Our method, sensGAN, estimates the worst-case latent confounder under constrained confounding strengths using an adversarial framework and provides a sensitivity contour diagnostic plot to suggest more reliable gene targets based on measured variables. In the following, we motivate the critical components of our framework one at a time: (1) quantifying the two “knobs” that define confounding strength, (2) defining the generative adversarial network (GAN) architecture that enables learning the confounder, and (3) training the GAN architecture.

### 2.1. Predictive gain: defining the confounding strength

We start with our formalization of quantifying the strength of a hypothetical latent confounder, which we denote as 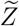 . Intuitively, the strongest latent confounder is the one that improves predictive performance the most for either the genes (*Y* ; “outcomes”) or the case-control status (*D*; treatment). Given 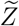, we can learn 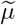 and 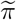 following the generative model in Eq. (1). The predictive gain of 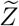 is defined as the normalized deviance-based partial *R*^2^ (DBPR) defined below, which we denote as *R*^2^.

For each prediction task (treatment *D* and outcomes *Y*), we compare three fitted models: (i) an unadjusted model that does not consider any latent confounders (“un”, see (2), (ii) an adjusted model using the *strongest* unobserved confounder 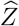 (the *k*-dimensional confounder that maximizes the likelihood), and (iii) an adjusted model using a particular hypothetical confounder 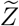. Let *ℓ*_*D*_(*π*) and *ℓ*_*Y*_ (*µ, α*) denote the Bernoulli and negative-binomial (NB) log-likelihoods under (1), and let *ℓ*_*D*,sat_ and *ℓ*_*Y*,sat_ denote the corresponding *saturated* (perfect-fit) log-likelihoods, i.e., the maximum achievable log-likelihood when each observation has its own free parameter. The deviance of a fitted model is Dev = 2(*ℓ*_sat_ − *ℓ*), with smaller deviance meaning a better fit. Dev_*D*_ is defined as the sum of Bernoulli deviances across all donors, and 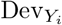 is defined as the sum of NB deviances across all genes. Details and derivations provided in Appendix S3. The deviance-based partial *R*^2^ for including *Z* into the model is the fraction of unadjusted deviance removed:

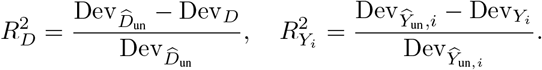

Finally, we normalize by the maximal gain achieved by 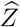 to obtain bounded predictive-gain “knobs” *κ, η* ∈ [0, 1], defined as

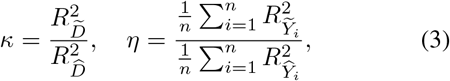

which quantify how predictive 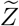 is relative to the strongest confounder in our model class. These emulate similar properties and interpretations of partial *R*^2^ for linear models as discussed in sensmarker (Cinelli & Hazlett, 2020). The normalized predictive gain denotes the proportional predictive gain introduced by an arbitrary 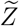, normalized by the maximal predictive gain introduced by the most powerful 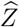 . A normalized predictive gain of 1 means that 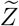 achieves 100% of the maximal predictive gain. Importantly, it is also differentiable, allowing gradient calculation in a deep learning model, as we describe later in this section.

### 2.2. Architecture and loss

Now that we have a quantification of the impact of a latent confounder, we next need to describe sensGAN’s deep-learning architecture, which allows us to learn the confounder *Z* that meets our specifications. Broadly speaking, sensGAN’s architecture has two fundamental components: the predictor and the generator. Here, we first describe each component of the architecture as shown in Fig. 3.

**Figure 3.**
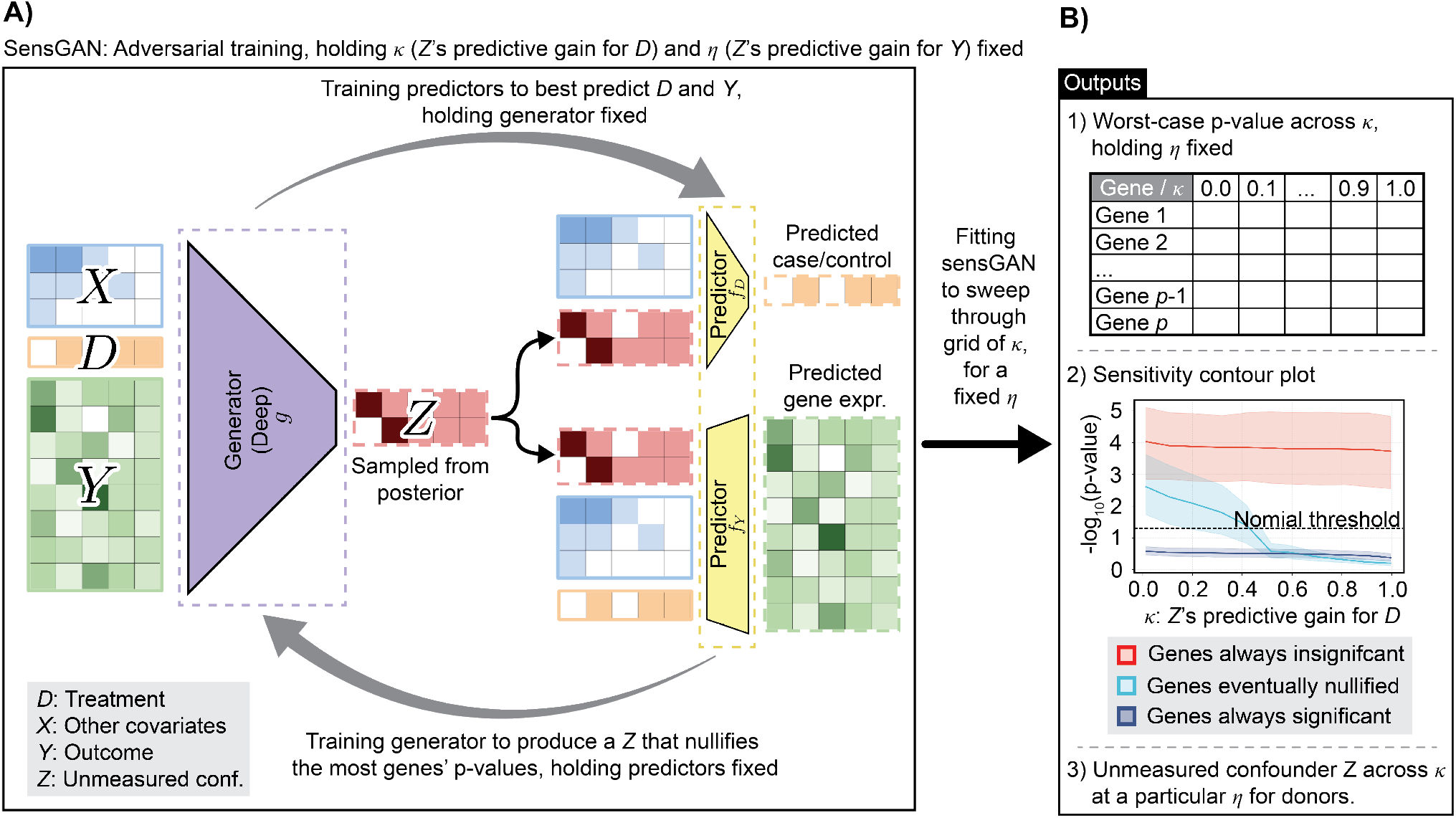
sensGAN method overview. **A)** The GAN system that learns the confounder *Z* with target predictive gains (*κ, η*), achieving predictive gains (*κ*^*′*^, *η*^*′*^). **B)** Method outputs while sweeping through a grid of *κ* and holding *η* fixed.

#### Predictors

The predictors *f*_*D*_(·) and *f*_*Y*_ (·) are 1-layer (i.e., shallow) fully connected networks that estimate model parameters by maximizing the likelihoods. To *D*_*i*_, the treatment predictor *f*_*D*_({*X*_*i*_, *Z*_*i*_}) uses a binary cross-entropy (BCE) loss. To predict *Y*_*i*_, the outcome predictor *f*_*Y*_ ({*D*_*i*_, *X*_*i*_, *Z*_*i*_}) uses a NB log-likelihood loss. Both predictors are designed to emulate logistic regression and NB GLM regression, respectively. There are no activation functions involved, so the weight corresponding to variable *D* in *f*_*Y*_ (·) can be interpreted as the “coefficient” *T*_*Y*_ in our generative model; see Eq. (1). The overdispersion parameters for modeling *Y* are estimated from the unadjusted NB model fitting.

#### Generator

The generator *g*(·) deterministically maps donor-level information to *k* latent confounders 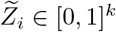 for donor *i*, based on covariates *X*_*i*_, treatment status *D*_*i*_, and outcome (gene expression) vector *Y*_*i*_. By default, the generator is a two-layer fully connected neural network with ReLU activation that produces 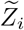 via a sigmoid output layer.

### 2.3. Training

Equipped with the predicator and generator architectures, we are now ready to explain how we train sensGAN. Drawing inspiration from the OVB literature (Cinelli & Hazlett, 2020; Veitch & Zaveri, 2020), we strive to learn latent con-founders 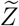 that nullifies as many genes (i.e., eliminates the significance of the largest number of genes associated with *D*), such that 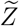 has a particular pre-specified predictive gain {*κ, η*}. The overall procedure has two main steps:

- **Step 1**: Learn the unadjusted model and the most powerful confounder, both of which are needed to define the normalized DBPRs; see Eq. (3),
- **Step 2**: Train the predictors *f*_*D*_(·) and *f*_*Y*_ (·) and generator *g*(·) in an adversarial manner.

We highlight each step below. The full pseudocode is described in Appendix S7.

#### Step 1: Unadjusted predictors and most powerful confounder

To operationalize our definition of the predictive gains in Eq. (3), we need to first compute the likelihood of the unadjusted model and the adjusted model using the most predictive confounder 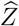 .

To learn the unadjusted models, we fit a logistic regression to predict *D* from *X*, and fit *p* NB GLMs to predict each gene in *Y* from *X* and *D*. From these fitted models, we can compute 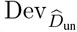 and 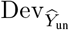, respectively. The gene-specific overdispersion parameters *α* estimated at this stage are held fixed and used throughout all subsequent sensGAN calculations.

We then train the predictors and the generator to minimize the joint negative log-likelihood, yielding the most powerful confounder *Z* and the corresponding models *f*_*D*_(·), *f*_*Y*_ (·), and *g*(·). Specifically, we define the BCE loss based on predicted logits as ℒ_BCE_ and the NB loss based on the predicted mean as ℒ_NB_. Predictors are trained respectively by minimizing ℒ_BCE_ and ℒ_NB_. Generators are trained by minimizing

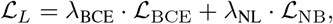

where *λ*_BCE_, *λ*_NL_ ≥ 0 are hyperparameters controlling the weight of their respective terms. From this fitted model, we can compute Dev_*D*_ and Dev_*Y*_, respectively. More details are discussed in Appendix S5.

#### Step 2: Adversarial training

We now describe how we train predictors *f*_*D*_(·) and *f*_*Y*_ (·) and generator *g*(·). After initializing the GAN architecture (see Appendix S6), we then update the predictors and generator alternatively and adversarially – the predictors maximize log-likelihoods while the generator controls for predictive gains. Specifically, in one iteration, the generator is frozen and the predictors *f*_*D*_(·) and *f*_*Y*_ (·) are updated; in the other, the predictors are frozen and the generator *g*(·) is updated.

When updating the predictors, we minimize ℒ_BCE_ to update the weights in *f*_*D*_(·) and minimize ℒ_NB_ to update the weights in *f*_*Y*_ (·). This is relatively straightforward, as both predictors are designed to emulate logistic and NB GLMs, respectively, where the optimization is convex.

Generator optimization is more involved and represents a key component of the sensGAN framework. The loss function when optimizing *g*(·) involves three main terms: the plausibility objective ℒ_P_, the significance objective ℒ_S_, and a regularizer ℒ_reg_. We explain each component below.

- To motivate the plausibility objective, we note that the strongest confounder 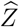 explains away all unexplained residuals in *D* and *Y*, by construction. This is unlikely to reflect any practical latent confounder. We thus define the *plausibility objective* (ℒ_P_): the predictive gains of plausible confounders 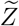 with respect to *D* and *Y* are constrained by prespecified targets, (*κ, η*) ∈ [0, 1],

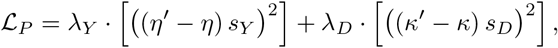

where *κ*^*′*^ and *η*^*′*^ denote the achieved predictive gains with respect to *D* and *Y*, respectively; *s*_*Y*_ and *s*_*D*_ are scaling factors, and *λ*_*Y*_, *λ*_*D*_ ≥ 0 are hyperparameters.
- To motivate the significance objective, we formalize “worst-case confounder” given *κ* and *η* as the confounder 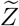 that nullifies the largest number of significant genes. Thus, we define the *significance objective* (ℒ_*S*_) that minimizes the average sigmoid-transformed excess of the test statistics over the significance threshold *δ* ∈ (0, 1) (i.e., typically *δ* = 0.05),

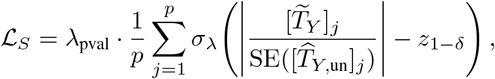

where 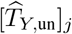 and 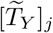 denote the estimated coefficients of *D* for gene *j* in the unadjusted and adjusted NB models, see Eq. (2) and (1) respectively. 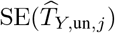 is the corresponding standard error from the unadjusted model. To compute the test statistic, we approximate 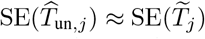 assuming comparable uncertainty before and after adjustment. *z*_1−*δ*_ is the (1 − *δ*) quantile of the standard normal distribution. *σ*_*λ*_(·) denotes a sigmoid function, which smoothly maps test statistics onto the unit interval. *λ*_pval_ ≥ 0 is a hyperparameter controlling the overall strength of the significance penalty.
- The last term is the *regularizer* (ℒ_reg_). This includes a correlation regularizer, a cosine similarity loss, and a variance-matching penalty, all of which we have empirically found help stabilize the learning of the latent confounder *Z* and prevent the model from collapsing to a constant value. See more details in Appendix S5.

Combining all the loss terms together, the objective function of the generator *g*(·) given pre-specified (and fixed) predictive gains *κ*^*^ and *η*^*^ is to minimize,

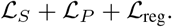

To span all plausible confounders, we train the sensGAN model sequentially by sweeping over the predictive gains (*κ, η*) on the unit interval [0, 1]. At the end of the sequential training, the generator finds a series of worst-case confounders with varied predictive gains.

The latent confounders *Z* can be used in a standard NB GLM analysis where we regress {*X, D, Z*} onto *Y*, where we record the significance of each gene’s coefficient in *T*_*Y*_, see Eq. (1). Gene-level p-values are computed using Wald statistics, where the test statistic for each gene is formed by dividing the adjusted coefficient by its standard error estimated from the unadjusted model 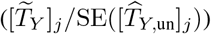. Mathematically, the p-value of each gene changes in the model that adjusts for our learned latent confounder, compared to the unadjusted model, see Eq. (2)). We are interested in the specific value of *κ*^*^ that “nullifies” the gene given a fixed *η*, if it is nullified in the adjusted model.

Additional details, such as the choice of hyperparameters, are deferred to Appendix S6. Given the potential instability of GAN training as mentioned by anonymous reviewers, we also discuss an alternative training strategy using bilevel optimization, where we elaborate on it in Appendix S6.3. Furthermore, we elaborate further on the modeling strategy, explaining why we chose to model scRNA-seq data as pseudobulked samples rather individual cells, as done in work like causarray (Du et al., 2026) and CoCoA-diff (Park & Kellis, 2021), in Appendix S4.

### 2.4. Downstream analysis: Calibration of predictive gains with measured covariates

From the sensitivity analysis, sensGAN produces a series of plausible worst-case confounders indexed by normalized predictive gains (*κ, η*) with respect to the exposure *D* and outcome *Y* . To interpret these latent confounders relative to observed covariates, we perform a calibration step that computes predictive gains for measured covariates, placing latent and measured effects on a common scale. If a worst-case confounder with strength *x* times that of the strongest measured covariate is required to nullify a gene’s significance, large values of *x* indicate that such confounding is potentially implausible, supporting the robustness of the discovery. See more details in Appendix S6.

### 2.5. Downstream analysis: Diagnostic sensitivity contour plots

To connect sensGAN’s core idea to gene-level robustness, we introduce a *sensitivity diagnostic* inspired by (Cinelli & Hazlett, 2020). Fixing the predictive-gain knob for the outcome (*η*) to reflect a conservative but bounded amount of confounding in the outcome, we vary the treatment-side knob (*κ*) and record the corresponding worst-case p-values. This assesses how predictive a latent confounder needs to be of *D* in order to nullify a gene’s significance. Genes that lose significance at small *κ* are sensitive to weak confounding, whereas genes that remain significant until large *κ* exhibit greater robustness.

## 3. Experimental Setup

### 3.1. Analysis of the simulation dataset for method comparison

We evaluate sensGAN on simulated data generated from a model with a single unobserved binary confounder (*n*=100, *p*=100, *d*=4, *k*=1). Observed covariates *X* and the binary latent confounder *Z* jointly influence a case–control variable *D* through a logistic regression model, while the outcome matrix *Y* is generated from a NB generalized linear model depending on *X, D*, and *Z*. By varying the effects of *D* and *Z* on gene expression, we simulate three groups of genes: (i) genes truly associated with *D* but not *Z*, (ii) genes associated with *Z* but not *D*, and (iii) genes associated with neither. More details and parameter settings are provided in Appendix S8.

To evaluate different methods, we first compare the correlations between the learned confounder and the true confounder (Fig. 4). The most powerful confounder identified by sensGAN shows a correlation comparable to that of causarray (Du et al., 2026) and RUVr (Risso et al., 2014), two surrogate-variable methods that we benchmark against. As the predictive-gain knob, *κ*, is turned from 1 to 0, the correlation between the learned and true confounders decreases smoothly from approximately 0.75 to 0.4. sensGAN not only identifies the strongest latent confounding structure, on par with competing approaches, but also generates a continuum of plausible worst-case confounders constrained by predictive gains.

**Figure 4.**
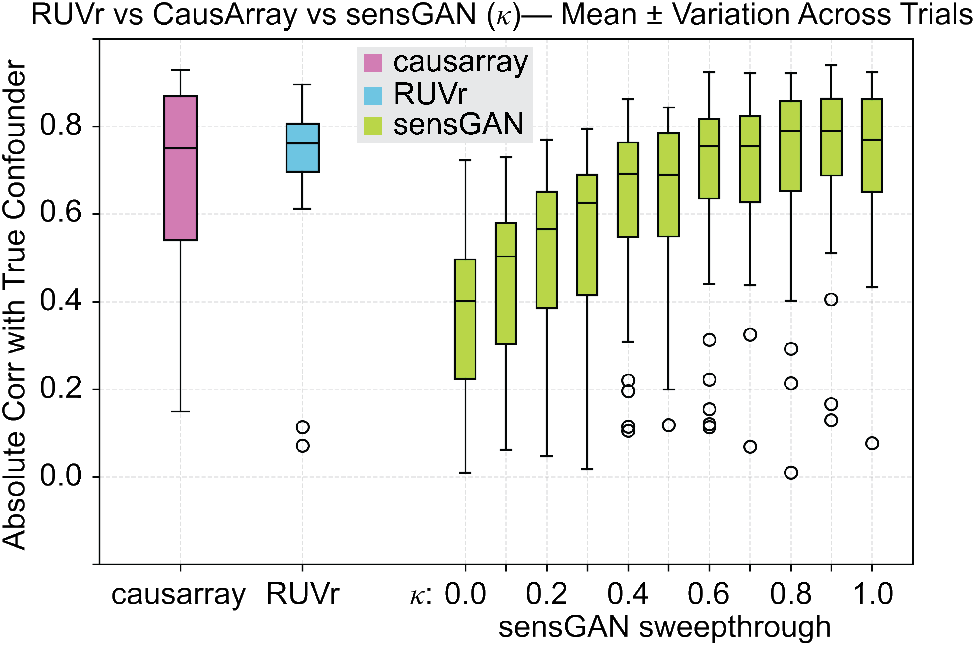
Correlation comparisons among methods. Correlations between the true confounder and the confounders estimated by causarray, RUVr, and sensGAN.

Next, we compare contingency tables of gene significance across methods (Table 1). All methods correctly identify always-significant genes, while causarray behaves conservatively for always-insignificant genes, likely due to its design for single-cell rather than donor-level pseudobulk data. Differences are most pronounced for nullified genes: the unadjusted GLM naively labels all such genes as significant (Fig. 4B), and causarray and RUVr split them between significant and insignificant categories. In contrast, sensGAN learns a family of plausible confounders with varying predictive gains and identifies a substantial fraction of nullified genes under worst-case confounding (8 out of 15; Table 1), while maintaining high accuracy for always-significant and always-insignificant genes (Fig. 5).

**Table 1.**
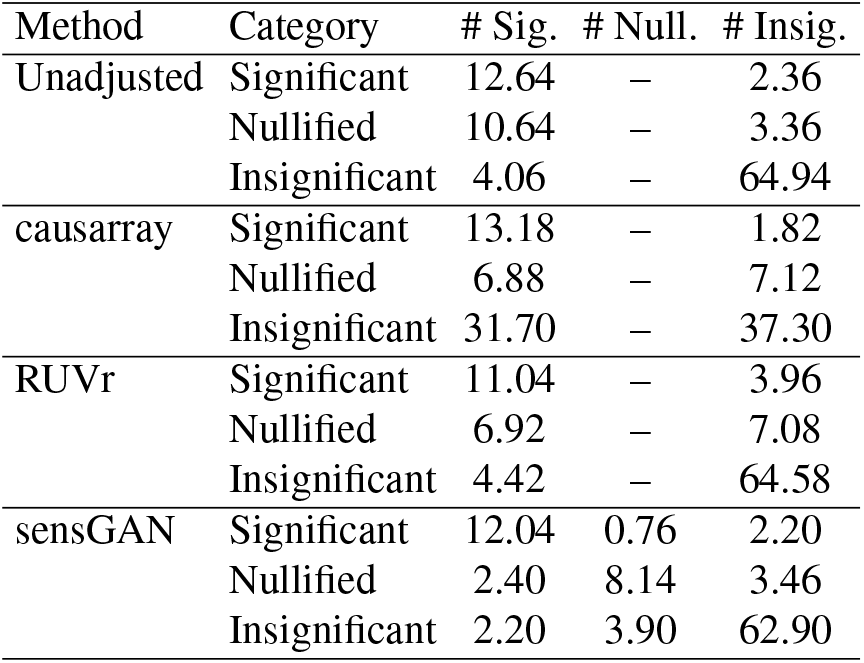
Comparison of mean gene counts across simulated significance categories. Rows: true gene categories, columns: inferred categories assigned by each method; values report mean counts averaged over 10 trials.

| Method | Category | # Sig. | # Null. | # Insig. |
| --- | --- | --- | --- | --- |
| Unadjusted | Significant | 12.64 | – | 2.36 |
|  | Nullified | 10.64 | – | 3.36 |
|  | Insignificant | 4.06 | – | 64.94 |
| causarray | Significant | 13.18 | – | 1.82 |
|  | Nullified | 6.88 | – | 7.12 |
|  | Insignificant | 31.70 | – | 37.30 |
| RUVr | Significant | 11.04 | – | 3.96 |
|  | Nullified | 6.92 | – | 7.08 |
|  | Insignificant | 4.42 | – | 64.58 |
| sensGAN | Significant | 12.04 | 0.76 | 2.20 |
|  | Nullified | 2.40 | 8.14 | 3.46 |
|  | Insignificant | 2.20 | 3.90 | 62.90 |

**Figure 5.**
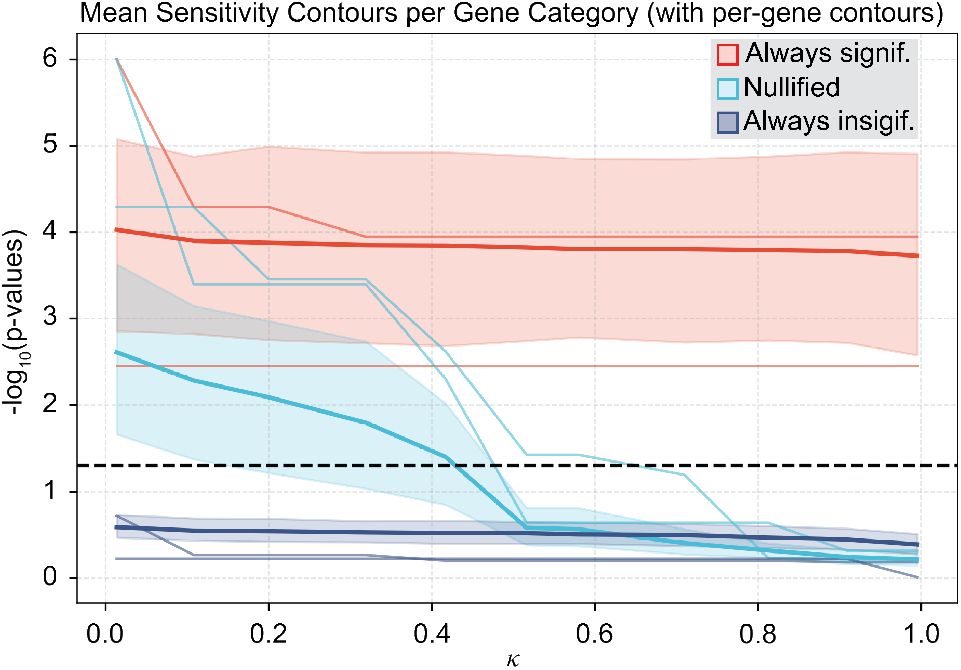
Simulation demonstrates sensGAN’s accuracy. Sensitivity diagnostic contour plot. With fixed *η* = 0.5, the thick contour curve represents the mean across genes, the thin curve shows an individual gene for illustration, and the shaded band denotes the 95% confidence interval.

We further present the sensitivity diagnostic contour plot based on sensGAN outputs (Fig. 4). We focus on nominal p-values rather than multiple-testing-adjusted p-values, as these allow us to better investigate how p-values change as the level of confounding increases. The y-axis is the logarithm of the worst-case p-values, the black dashed line is the significance threshold of 0.05, and the x-axis is the predictive knob (*κ*) for the treatment. Every gene has a contour curve that depicts its sensitivity to the estimated latent confounder, exemplified by the thin curves. The thick curves are the mean contour curves, and the bands are the 95% confidence intervals, colored by the simulated gene categories. If a gene crosses the significance threshold with a smaller *κ*, the gene is considered to be nullified by a weaker confounder. Compared to genes nullified by stronger confounders, this gene is less directly associated with the treatment. Therefore, the differential expression conclusion is less robust.

### 3.2. Validation of contour curve accuracy

Having shown that sensGAN recovers latent confounders correlated with those identified by existing methods, we next evaluate its ability to accurately characterize sensitivity contour curves, which prior approaches do not provide. Our primary focus is to investigate whether sensGAN can faithfully track how gene-level p-values evolve with increasing confounding strength. To this end, we simulate data as before (*n* = 100, *p* = 104, *d* = 4, *k* = 1), but consider a continuous confounder *Z* to enable finer control over predictive gains, and restrict attention to genes that are associated with *Z* but not with *D* (Category (ii)). We first construct reference sensitivity contour curves using a crawler-based search that manually explores the predictive-gain space to identify the “true” worst-case confounder for a particular setting of (*κ, η*) (see Appendix S8). We then apply sensGAN to the same simulated data and compare the resulting sensitivity contours, assessing whether sensGAN accurately reproduces the true nullification behavior across predictive-gain levels.

We first evaluate sensGAN at the gene level by comparing per-gene sensitivity contour curves against the reference contours (Fig. 5), with a primary focus on the difference in *κ*^*^ at the nullification point. Quantitatively, the difference is modest, with a mean difference of 0.05 and a 95% confidence interval of (0.00, 0.22), indicating close alignment in the predictive-gain level at which genes lose significance.

We next compare the mean sensitivity contour curves across genes (Fig. 6B). Overall, the mean curves produced by sensGAN closely track the reference contours, with similar shapes and comparable crossing points near *κ*≈ 0.95. The 95% confidence intervals largely overlap across the predictive-gain range, suggesting that sensGAN captures the dominant trend in how significance decays with increasing confounding strength. See Appendix S9 for more results.

**Figure 6.**
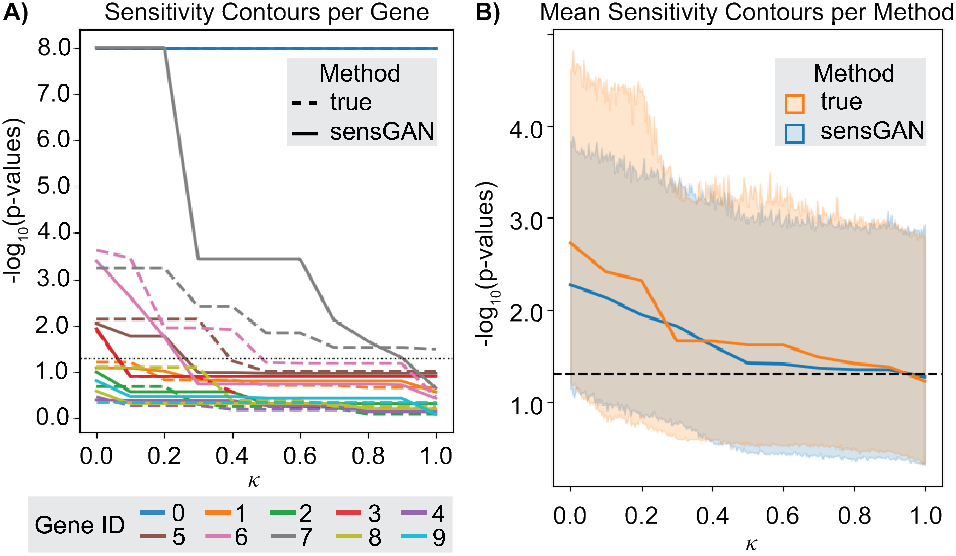
Sensitivity contour plot comparison between the reference and the sensGAN output. **A)** Per-gene sensitivity contour plots showing − log_10_(*p*) as a function of the predictive-gain parameter *κ* for individual genes. Dashed curves denote reference contours obtained via the crawler, while solid curves denote contours recovered by sensGAN. **B)** Mean sensitivity contour curves across genes, with shaded regions indicating 95% confidence intervals.

### 3.3. Analysis of microglia studying Alzheimer’s disease, adjusting for unmeasured co-occurring neurodegenerative disease

To demonstrate how sensGAN can clarify the biological signals, we showcase our analysis of systemic lupus erythematosus (SLE) and Alzheimer’s disease (AD). We discuss the AD case-study here in detail, while deferring the SLE case-study to Appendix S10. For our AD case-study, we leverage the Seattle Alzheimer’s Disease Brain Cell Atlas (SEA-AD) consortium (Gabitto et al., 2024) and specifically focus on the microglia in the prefrontal cortex (PFC) due to its critical role in clearing AD pathology and its implication from established GWAS studies (Karch & Goate, 2015; Scheltens et al., 2016).

A major challenge in studying cellular changes during AD progression is that post-mortem donors often exhibit multiple neurodegenerative diseases beyond AD itself (Nichols et al., 2023; McAleese et al., 2021). These co-occurring diseases (“co-pathologies”) can confound single-cell analyses. For example, only 31% of late-onset AD cases show AD-specific signatures (Robinson et al., 2023).

We construct a pseudobulk gene expression matrix per donor (i.e., *Y*) and perform our sensGAN analysis to determine which genes are impacted based on whether the donor has AD pathology after post-mortem dissection of their brain tissue (i.e., *D*), adjusting for sex, age, APOE4 status, ethnicity, and post-mortem interval (PMI) (i.e., *X*). Since we are only interested in whether or not significant genes remain significant during our sensitivity analysis, we focus on a key set of genes that have a p-value *<* 0.2 when doing a differential expression test via DESeq2 (Love et al., 2014), resulting in *p* = 448 genes in our analysis. Our goal is to determine which nominally significant genes are nullified by the latent confounders identified by sensGAN (Goal #1) and what these latent confounders represent biologically (Goal #2).

Our sensGAN analysis, sweeping across *κ* from 0 to 1, disentangles potential microglial pathways that may be confounded by co-pathologies from those that are specific to AD. Our diagnostic contour plots reveal that genes that are always nominally significant are enriched for an apoptosis pathway (red; Fig. 7) (Dou et al., 2024). For instance, the *LGALS3* gene is strongly upregulated in plaque-associated microglia in AD models and human tissue (Tan et al., 2021), and has been implicated to be AD-specific through knock-outs in AD mouse models, which demonstrated reduced plaque burden and improved cognitive behaviors (Siew et al., 2024). In stark contrast, genes that are eventually nullified by sensGAN’s learned latent confounders *Z* are enriched for transmembrane transport (light blue; Figure 7). For instance, AHNAK regulates voltage-gated calcium channels (Matza et al., 2008). Although it is nominally significant without confounder adjustment, this gene is quickly nullified by sensGAN. This is plausible, since its association with AD is not solely through microglia but also through its interaction with neurons (Wang et al., 2025). Furthermore, it has also been suggested to be involved in Lewy bodies (Santpere et al., 2018) and Frontal Temporal Dementia (Lorenzini et al., 2023), two other co-occurring pathologies typically associated with AD. Additionally, AQP9, a gene that regulates membrane channels to conduct water, is also nullified, which is canonically associated with an AD-relevant gene for astrocytes, not microglia (Liu et al., 2018). Together, these results demonstrate that sensGAN can meaningfully “purify” DEG results, retaining only microglia-specific DEGs robust to non-AD latent confounders.

**Figure 7.**
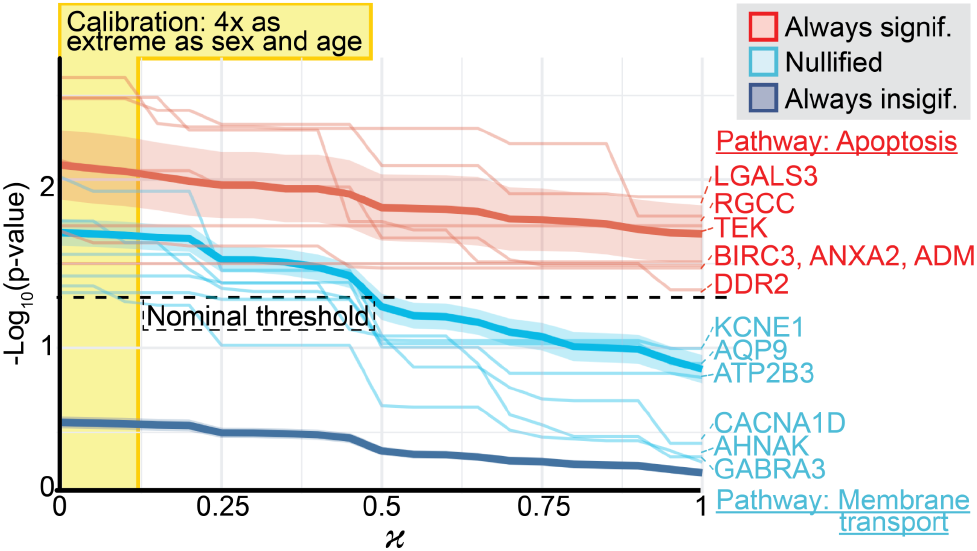
sensGAN prioritizes DEGs specific to microglia in AD. Diagnostic contour, showing genes that are always significant (red), nullified (light blue), and always insignificant (dark blue) across different values of *κ*. The calibration with respect to age and sex is shown (yellow).

Next, we illustrate that sensGAN’s latent confounders are suggestive of currently measured co-occurring neurodegenerative disease. While we do not use any co-pathologies in our sensGAN analysis, the SEA-AD consortium also stages all the post-mortem tissues for arteriolosclerosis (i.e., a vascular disease), Lewy bodies, and Limbic-predominant age-related TDP-43 encephalopathy (LATE). We compute how well sensGAN’s learned confounders predict these co-pathologies (Fig. 8). Surprisingly, for *κ* ∈ [0.5, 1], the predictive power measured by *R*^2^ between the predicted (linear regression) and observed co-pathology stages is significantly non-zero and not changing. This suggests that sensGAN latent confounders are capturing co-pathology impacts from snRNA-seq data, even with a modest value of *κ* = 0.5. We hypothesize that the latent confounders estimated by sensGAN additionally capture effects beyond co-pathology, for instance, the effects of spatial microenvironments, genetics, or environmental factors (e.g., diet) on the donor that affect AD.

**Figure 8.**
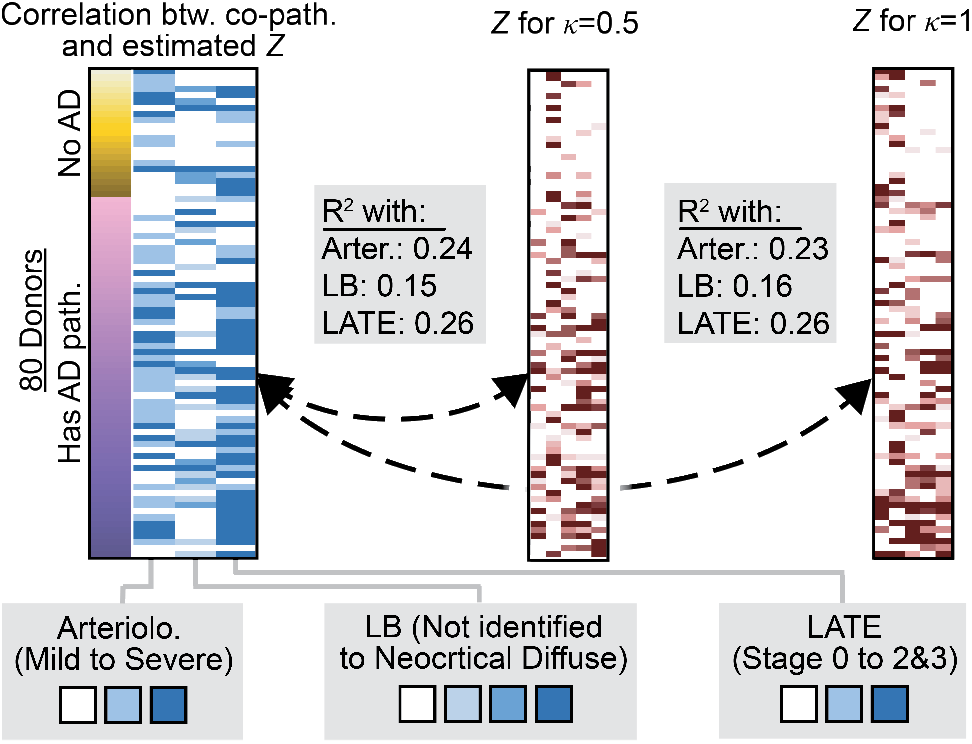
sensGAN’s latent confounders reflect co-pathologies. Donors (rows) and their measured co-pathology, as well as the estimated latent confounders for *κ* ∈ {0.5, 1}.

We further examine which microglial states harbor the DEGs identified by sensGAN. Although sensGAN operates on pseudobulk data, mapping DEGs to microglial states provides biological context (Prater et al., 2023). We first visualize the microglia in the SEA-AD cohort by their annotated state (Fig. 9A). Significant genes that are either never nullified by sensGAN (*LGALS3, ANXA2*) or eventually nullified (*AQP9, AHNAK*) show elevated expression primarily in DAM and transitory microglia (Fig. 9B). These results highlight how sensGAN can prioritize DEGs based on their robustness to latent confounding while preserving biologically meaningful microglial localization.

**Figure 9.**
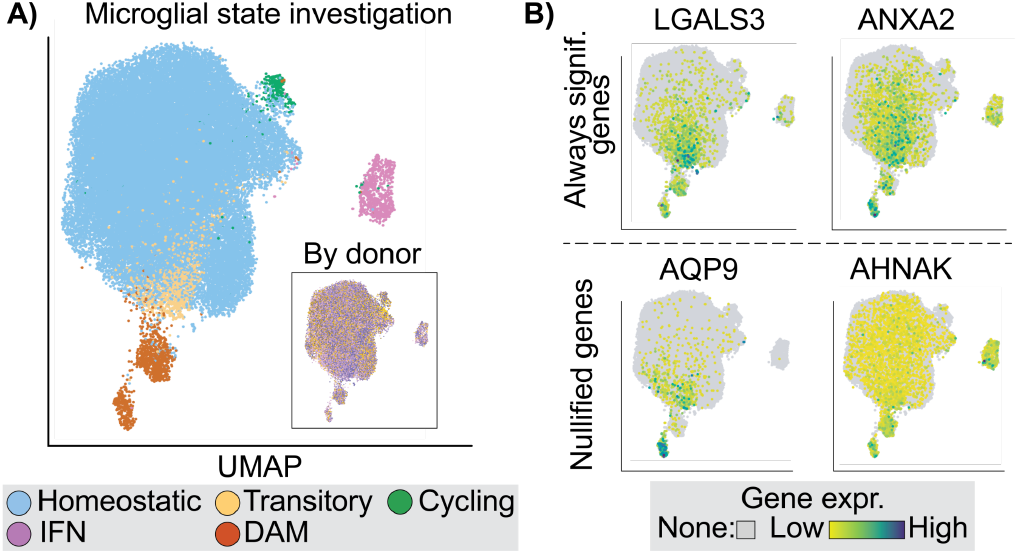
sensGAN prioritizes DEGs specific to microglia in AD. **A)** UMAP of single-nuclei microglia in SEA-AD, colored by microglial state and donor. **B)** Expression of certain genes across the microglia.

### 3.4. Discussion

We present sensGAN, an adversarial framework that shifts the focus of high-dimensional genomic analysis from binary significance to quantitative robustness. Unlike traditional surrogate-variable methods that provide single-point estimates of latent factors, sensGAN explicitly explores the confounding spectrum by learning “worst-case” latent variables under controlled predictive-gain constraints. Our simulations demonstrate that sensGAN maintains high fidelity in separating robust biological signals from those easily nullified by hidden confounders, effectively capturing a continuum of plausible confounding scenarios that point-estimation methods often miss. By treating confounding as a formal machine learning optimization problem, sensGAN provides a principled metric for determining how strong a latent process must be to explain away a discovery.

Applied to complex diseases, sensGAN successfully “purifies” differential expression results, isolating pathways specifically associated with AD from signals likely driven by co-occurring neurodegenerative disease. For instance, our framework prioritizes apoptosis-related processes as robust AD-specific drivers, while identifying transmembrane transport genes as highly sensitive to unmeasured co-occurring neurodegenerative disease, such as Lewy body pathology. While currently limited by its reliance on pseudobulk aggregation and calibrated (rather than absolute) effect sizes, sensGAN establishes a unified, future-oriented approach to interpreting single-cell genomics in the presence of pervasive, evolving confounding.

## Acknowledgments

We thank Suman Jayadev, Eardi Lila, Duolan Ouyang, Katherine E. Prater, Aquene N. Reid, Taek Son, the anonymous ICML reviewers, and other members of the Lin Lab for support, helpful discussions, and feedback on this work. Research reported in this manuscript was supported by the University of Washington’s Royalty Research Fund (RRF) and the National Institute of General Medical Sciences (NIGMS) of the National Institutes of Health (NIH) under award number: R35GM162089.

The results relevant to SEA-AD published here are based on data obtained from the The AD Knowledge Portal DOI: https://doi.org/10.7303/9618137. Study data were generated from postmortem brain tissue obtained from the University of Washington BioRepository and Integrated Neuropathology (BRaIN) laboratory and Precision Neuropathology Core, which is supported by the NIH grants for the UW Alzheimer’s Disease Research Center (P50AG005136 and P30AG066509) and the Adult Changes in Thought Study (U01AG006781 and U19AG066567). This study is supported by NIA grant U19AG060909.

The NACC database is funded by NIA/NIH Grant U24 AG072122. NACC data are contributed by the NIA-funded ADRCs: P30 AG062429 (PI James Brewer, MD, PhD), P30 AG066468 (PI Oscar Lopez, MD), P30 AG062421 (PI Teresa Gomez-Isla, MD), P30 AG066509 (PI Thomas Grabowski, MD), P30 AG066514 (PI Mary Sano, PhD), P30 AG066530 (PI Helena Chui, MD, Arthur Toga, PhD), P30 AG066507 (PI Marilyn Albert, PhD), P30 AG066444 (PI David Holtzman, MD), P30 AG066518 (PIs Lisa Silbert, MD, Kevin Duff, PhD), P30 AG066512 (PI Thomas Wisniewski, MD), P30 AG066462 (PI Scott Small, MD), P30 AG072979 (PI David Wolk, MD), P30 AG072972 (PIs Charles DeCarli, MD, Rachel Whitmer, PhD), P30 AG072976 (PI Andrew Saykin, PsyD), P30 AG072975 (PI Julie Schneider, MD, MS), P30 AG072978 (PI Ann McKee, MD), P30 AG072977 (PI Robert Vassar, PhD), P30 AG066519 (PI Joshua Grill, PhD), P30 AG062677 (PIs Brad Boeve, MD, Ronald Petersen, MD, PhD), P30 AG079280 (PI Jessica Langbaum, PhD), P30 AG062422 (PI Gil Rabinovici, MD), P30 AG066511 (PI Allan Levey, MD, PhD), P30 AG072946 (PI Linda Van Eldik, PhD), P30 AG062715 (PI Sanjay Asthana, MD, FRCP), P30 AG072973 (PI Russell Swerdlow, MD), P30 AG066506 (PIs Glenn Smith, PhD, ABPP, David Lowenstein, PhD, Ranjan Duara, MD), P30 AG066508 (PIs Stephen Strittmatter, MD, PhD, Christopher Van Dyck, MD), P30 AG066515 (PI Victor Henderson, MD, MS), P30 AG072947 (PI Suzanne Craft, PhD), P30 AG072931 (PI Henry Paulson, MD, PhD), P30 AG066546 (PIs Sudha Seshadri, MD, Gladys Maestre, MD, PhD), P30 AG086401 (PI Erik Roberson, MD, PhD), P30 AG086404 (PI Gary Rosenberg, MD), P30 AG086403 (PI Angela Jef-ferson, PhD), P30 AG072958 (PIs Heather Whitson, MD, Gwenn Garden, MD, PhD), P30 AG072959 (PI Jagan Pillai, MD, PhD), P30 AG092752 (Ihab Hajjar, MD, MS).

## Impact statement

This work introduces an adversarial framework to quantify the robustness of high-dimensional genomic findings against latent confounding. By transitioning from binary significance to a quantitative exploration of the confounding spectrum, this method facilitates more reliable interpretation of biological mechanisms and the prioritization of robust therapeutic targets. This advancement enhances the reproducibility and trustworthiness of computational biology, with potential long-term societal benefits for precision medicine across diverse complex diseases.

## Notes

https://github.com/yifanlinz/AD_sensitivity_ICML

### Competing Interest Statement

The authors have declared no competing interest.

### Summary of Updates

Fixed broken cross-references between the main text and appendix.

