## Supplemental Materials for "High-Dimensional Sensitivity Analysis for Genomic Studies: An Adversarial Framework for Learning Worst-Case Latent Confounders"

#### S1 Code and data availability

The sensGAN method is publicly available at `https://github.com/yifanlinz/AD_sensitivity_ICML`, which includes both the implementation of the method and the way it is used on different datasets in this work.

The systemic lupus erythematosus data (Perez et al., 2022), which we analyze in Section S10, is publicly available and can be downloaded at `https://www.ncbi.nlm.nih.gov/geo/query/acc.cgi?acc=GSE174188`. The Alzheimer's dataset (Gabitto et al., 2024) is publicly available and can be downloaded from `https://registry.opendata.aws/allen-sea-ad-atlas/`, and we focus on analyzing the microglial data from the prefrontal cortex.

#### S2 Motivation for confounder modeling

The challenge of latent confounding is particularly acute in Alzheimer's Disease (AD) research, where donors frequently exhibit complex comorbidity patterns or co-pathologies. Current computational methods struggle to isolate "direct" disease signals from these co-occurring processes.

**Comparison with Omitted Variable Bias (OVB) framework.** Methods such as sensmaker (Cinelli & Hazlett, 2020) utilize partial $R^2$ to reparameterize bias in linear models. While foundational, these approaches are not easily generalized to the non-linear Negative Binomial (NB) distributions required for pseudobulked single-cell RNA-seq data. sensGAN addresses this gap by enabling a formal sensitivity analysis across thousands of concurrent outcomes (genes) within a deep-learning adversarial framework.

**Comparison with Surrogate Variable Analysis (SVA).** Standard genomic adjustment tools, including RUVr (Risso et al., 2014), CATE (Wang et al., 2017), GCATE (Du et al., 2025), and causarray (Du et al., 2026), focus on capturing residual variation through low-dimensional surrogates. While these provide explicit latent representations, they typically treat confounding as a fixed correction rather than a spectrum. Because these estimates are often independent of the treatment variable, they may fail to reflect biologically relevant latent factors—such as vascular disease or Lewy body pathology—that are statistically associated with both the AD case-control status and the observed gene expression matrix.

#### S3 Details on deviance, DBPR, and normalized predictive gains

In Section S3.1, we describe the four different models used in the calculation of the DBPR. In Sections S3.2 and S3.3, we formally define deviance and DBPR. In Section S3.4, we formally define the "knobs" $\kappa$ and $\eta$ used in sensGAN to quantify the predictive gain of a latent confounder.

#### S3.1 Model family and fitted quantities.

For donors $i \in \{1, \dots, n\}$, let $D_i \in \{0, 1\}$ denote treatment (i.e., case-control status), $X_i \in \mathbb{R}^d$ observed covariates, and $Y_{ij}$ the (pseudobulk) count for gene $j \in \{1, \dots, p\}$. For any candidate latent confounder matrix $Z \in \mathbb{R}^{n\times k}$, our generative family induces (i) a Bernoulli model for $D$ with fitted linear predictor $\pi$ (notation as in the main text), and (ii) a negative-binomial (NB) model for $Y$ with fitted mean $\mu$ and gene-specific dispersion $\alpha$. We write the corresponding log-likelihoods as $\ell_D(\pi)$ and $\ell_Y(\mu, \alpha)$.

We instantiate the models in four standard ways for each task:

1. **Saturated model (sat)**: The saturated models are the *perfect-fit* references that achieve the maximum possible log-likelihood. Denote these maxima by $\ell_{D,\text{sat}}$ and $\ell_{Y,\text{sat}}$.
2. **Unadjusted model (un)**: Fit models *without* $Z$ to obtain $(\widehat{\pi}_{\text{un}}, \widehat{\mu}_{\text{un}}, \widehat{\alpha}_{\text{un}})$ and log-likelihoods $\ell_D(\widehat{\pi}_{\text{un}})$, $\ell_Y(\widehat{\mu}_{\text{un}}, \widehat{\alpha}_{\text{un}})$. In the following models, overdispersion is fixed at $\widehat{\alpha}_{\text{un}}$.
3. **Most predictive confounder (Denoted by $\widehat{\cdot}$ )**: Define $\widehat{Z}$ as a confounder of dimension-$k$ that yields the best achievable fit when included in the models with unadjusted overdispersion $\alpha$, i.e., it maximizes the likelihood (equivalently minimizes deviance). After fitting, this produces $(\widehat{\pi}, \widehat{\mu})$ and the log-likelihoods $\ell_D(\widehat{\pi})$, $\ell_Y(\widehat{\mu}, \widehat{\alpha}_{\text{un}})$.
4. **Arbitrary confounder (Denoted by $\widetilde{\cdot}$ )**: For any specific confounder $\widetilde{Z}$ (e.g., produced by sensGAN under constraints), fit models with the unadjusted overdispersion $\alpha$ including $\widetilde{Z}$ to obtain $(\widetilde{\pi}, \widetilde{\mu})$ and the log-likelihoods $\ell_D(\widetilde{\pi})$, $\ell_Y(\widetilde{\mu}, \widehat{\alpha}_{\text{un}})$.

#### S3.2 Deviance

For modeling $D$, the Bernoulli deviance based on logits $\text{Dev}_D \in \mathbb{R}$ is defined:

$$\text{Dev}_{\widehat{D}_{\text{un}}} = \sum_{i=1}^{n} 2\big(\ell_{D_i,\text{sat}} - \ell_{D_i}(\widehat{\pi}_{\text{un}})\big), \quad \text{Dev}_{\widehat{D}} = \sum_{i=1}^{n} 2\big(\ell_{D_i,\text{sat}} - \ell_{D_i}(\widehat{\pi})\big), \quad \text{Dev}_{\widetilde{D}} = \sum_{i=1}^{n} 2\big(\ell_{D_i,\text{sat}} - \ell_{D_i}(\widetilde{\pi})\big).$$

For modeling $Y$, the NB deviance $\text{Dev}_Y = \{\text{Dev}_{Y_1}, \text{Dev}_{Y_2}, ..., \text{Dev}_{Y_n}\} \in \mathbb{R}^n$ is the sum across genes:

$$\begin{aligned}
\text{Dev}_{\widehat{Y}_{\text{un}},i} &= \sum_{j=1}^{p} 2\big(\ell_{Y_{ij}\text{sat}} - \ell_{Y_{ij}}(\widehat{\mu}_{ij,\text{un}}, \widehat{\alpha}_{j,\text{un}})\big), \\
\text{Dev}_{\widehat{Y}i} &= \sum_{j=1}^{p} 2\big(\ell_{Y_{ij},\text{sat}} - \ell_{Y_{ij}}(\widehat{\mu}_{ij}, \widehat{\alpha}_{j,\text{un}}\big), \\
\text{Dev}_{\widetilde{Y}i} &= \sum_{j=1}^{p} 2\big(\ell_{Y_{ij},\text{sat}} - \ell_{Y_{ij}}(\widetilde{\mu}_{ij}, \widehat{\alpha}_{j,\text{un}}\big).
\end{aligned}$$

Thus, deviance behaves like a likelihood-based error metric: lower deviance indicates a better fit.

#### S3.3 DBPR: deviance-based partial $R^2$

We quantify the *incremental predictive gain* ($R^2_D \in \mathbb{R}$, $R^2_Y = \{R^2_{Y_1}, R^2_{Y_2}, \dots, R^2_{Y_n}\} \in \mathbb{R}^n$) of adding a confounder $Z$ by the fraction of unadjusted deviance it removes.

Using the fitted models above, this yields the quantities shown in the main text:

$$\widehat{R}^2_D = \frac{\text{Dev}_{\widehat{D}_{\text{un}}} - \text{Dev}_{\widehat{D}}}{\text{Dev}_{\widehat{D}_{\text{un}}}}, \quad \widehat{R}^2_{Y_i} = \frac{\text{Dev}_{\widehat{Y}_{\text{un}},i} - \text{Dev}_{\widehat{Y}_i}}{\text{Dev}_{\widehat{Y}_{\text{un}},i}}, \qquad \widetilde{R}^2_D = \frac{\text{Dev}_{\widehat{D}_{\text{un}}} - \text{Dev}_{\widetilde{D}}}{\text{Dev}_{\widehat{D}_{\text{un}}}}, \quad \widetilde{R}^2_{Y_i} = \frac{\text{Dev}_{\widehat{Y}_{\text{un}},i} - \text{Dev}_{\widetilde{Y}_i}}{\text{Dev}_{\widehat{Y}_{\text{un}},i}}.$$

##### S3.4 Normalized DBPR (predictive-gain knobs)

Because $\widehat{Z}$ is defined to be the *most predictive* confounder in our model class, $\widehat{R}^2_D$ and $\widehat{R}^2_Y$ represent maximal attainable deviance reduction for predicting $D$ and $Y$, respectively. We therefore define normalized predictive gains for any $\widetilde{Z}$ by

$$\kappa = \frac{\widetilde{R}^2_D}{\widehat{R}^2_D}, \qquad \eta = \frac{\frac{1}{n}\sum_{i=1}^n \widetilde{R}^2_{Y_i}}{\frac{1}{n}\sum_{i=1}^n \widehat{R}^2_{Y_i}},$$

so that $\kappa, \eta \in [0, 1]$ quantify the *fraction of maximal predictive gain* achieved by $\widetilde{Z}$ for $D$ and $Y$.

#### S4 SenGAN architectures

In Section S4.1, we discuss why we find it better to model latent confounders at the donor level (via pseudobulking the gene expression vectors) rather than at the cell level. In Section S4.2, we define the generator architecture used in sensGAN to compute the latent confounder for each donor.

##### S4.1 Rationale for modeling scRNA-seq data on a donor level

We explain why we model sensGAN at the "donor level" (pseudobulking all cells from each of the $n$ donors, despite having scRNA-seq data) rather than at the "cell level." Specifically, this means we sum all the gene expression vectors of each cell to represent the gene expression vector for donor $i$, for $i \in \{1, \ldots, n\}$. Our choice to pseudobulk the data might seem counterintuitive, since there are existing methods such as causarray (Du et al., 2026) and CoCoA-diff (Park & Kellis, 2021) that model latent confounders at the cell level. However, we note some important distinctions that motivate our choice of pseudobulking the data, despite having scRNA-seq data:

- **Intended goal of adjusting for latent confounders that represent future neuropathologies**: As we mention in Section 1.1 in the main text, one of the primary motivations of sensGAN is to have the estimated latent confounders represent potential co-occurring neurodegenerative diseases. This is fundamentally a donor-level quantity. In contrast, methods such as causarray (Du et al., 2026) and CoCoA-diff (Park & Kellis, 2021) learn cell-level confounders, which do not scientifically represent the same goal as our intended goal.
- **Considerations for the complexity of sensGAN architecture and computational efficiency**: Consider the architecture of the generator, which we describe in Section S4.2. If we model the data at the cell level, we would need to adjust the architecture so that the donor-level confounder can be "broadcasted" to all cells from that donor. While this is feasible, it would dramatically increase the computational burden of fitting sensGAN and, hence, be detrimental to the goal of learning latent confounders.

##### S4.2 Generator architecture

The sigmoid-head generator directly maps the input to a bounded confounder via a deterministic neural network:

$$h = \phi(W_1[X, D, Y] + b_1),$$
$$\widetilde{Z} = \sigma(W_2 h + b_2),$$

where $\phi(\cdot)$ denotes a ReLU activation and $\sigma(\cdot)$ is the elementwise sigmoid function. This parameterization produces $\widetilde{Z}$ deterministically and is simple and stable to optimize. In practice, we initialize the final-layer weights $(W_2, b_2)$ by fitting a ridge regression in logit space, so that $\sigma(W_2 h + b_2) \approx Z^{(0)}$, where $Z^{(0)}$ is an initial confounder estimate.

### S5 Loss details

Here, we include more details about the explicit calculations of the loss terms for the different components of sensGAN. In Section S5.1, we define the likelihood loss used in the treatment predictor and the outcome predictor. In Section S5.2, we define all the terms of the regularization term $\mathcal{L}_{\text{reg}}$ used when training in the generator in the adversarial training step of sensGAN.

#### S5.1 Likelihood Loss

For the treatment predictor $f_D(\cdot)$, we use the binary cross-entropy loss with logits. Let $\widehat{s}_i \in \mathbb{R}$ denote the predicted logit for donor $i$, and let $\widehat{\pi}_i = \sigma(\widehat{s}_i)$ be the corresponding predicted treatment probability, where $\sigma(\cdot)$ denotes the logistic function. The per-donor loss is given by

$$\ell_i^{\text{BCE}} = \max(\widehat{s}_i, 0) - \widehat{s}_i D_i + \log\Big(1 + e^{-|\widehat{s}_i|}\Big),$$

which is a numerically stable implementation of binary cross-entropy with logits loss.

For the outcome predictor $f_Y(\cdot)$, we model gene expression counts with a negative binomial distribution. Let $Y_{ij}$ denote the count for donor $i$ and gene $j$, with mean $\mu_{ij} > 0$ and dispersion $\alpha_g > 0$. Define $r_j = 1/\alpha_j$. The per-entry negative log-likelihood is

$$\ell_{ij}^{\text{NB}} = -\Bigg[\log\Gamma(Y_{ij} + r_j) - \log\Gamma(r_j) - \log\Gamma(Y_{ij}+1) + r_j \log\Big(\tfrac{r_j}{r_j+\mu_{ij}}\Big) + Y_{ij}\log\Big(\tfrac{\mu_{ij}}{r_j+\mu_{ij}}\Big)\Bigg].$$

We aggregate across genes to obtain a per-donor loss,

$$\ell_i^{\text{NB}} = \sum_{j=1}^{p} \ell_{ij}^{\text{NB}},$$

.

When training the most powerful predictor, the loss function used to train both $f_D(\cdot)$ and $f_Y(\cdot)$ is,

$$\mathcal{L}_L = \lambda_{\text{BCE}} \cdot \mathcal{L}_{\text{BCE}} + \lambda_{\text{NB}} \cdot \mathcal{L}_{\text{NB}}, \quad \text{where} \quad \mathcal{L}_{\text{BCE}} = \frac{1}{n}\sum_{i=1}^{n} \ell_i^{\text{BCE}}, \quad \mathcal{L}_{\text{NB}} = \frac{1}{n}\sum_{i=1}^{n} \ell_i^{\text{NB}}.$$

#### S5.2 Regularization term

When training sensGAN via alternating adversarial optimization, we include a regularization term $\mathcal{L}_{\text{reg}}$ to stabilize generator updates and prevent degenerate confounder solutions. Let $\widetilde{Z} \in \mathbb{R}^{n\times k}$ denote the current generator output and $\widehat{Z} \in \mathbb{R}^{n\times k}$ the strongest confounder used as a reference. The regularization consists of the following components.

The correlation regularizer

$$\mathcal{L}_{\text{corr}}(\widetilde{Z}, \widehat{Z}) = \frac{1}{k}\sum_{a=1}^{k}\big(1 - \text{Corr}(\widetilde{Z}_{\cdot a}, \widehat{Z}_{\cdot a})\big)^2 + \frac{1}{k(k-1)}\sum_{a\neq b}\text{Corr}(\widetilde{Z}_{\cdot a}, \widehat{Z}_{\cdot b})^2,$$

encourages each generated confounder dimension to align with its corresponding reference dimension while discouraging cross-dimensional mixing.

The cosine similarity regularizer

$$\mathcal{L}_{\cos}(\widetilde{Z}, \widehat{Z}) = 1 - \frac{\langle \text{vec}(\widetilde{Z}), \text{vec}(\widehat{Z})\rangle}{\|\text{vec}(\widetilde{Z})\|_2 \, \|\text{vec}(\widehat{Z})\|_2},$$

promotes global alignment between the generated and reference confounders up to scale.

Table S1: Pearson correlation between estimated latent variable $\widehat{Z}$ and simulated $Z$ across $\kappa$ targets under different regularization settings.

| $\kappa$ Target | Full Reg. | No Var.-Matching Reg. (1) |
|---|---|---|
| 0.0 | 0.401 | 0.391 |
| 0.1 | 0.503 | 0.357 |
| 0.2 | 0.566 | 0.543 |
| 0.3 | 0.625 | 0.727 |
| 0.4 | 0.692 | 0.701 |
| 0.5 | 0.690 | 0.676 |
| 0.6 | 0.755 | 0.802 |
| 0.7 | 0.755 | 0.602 |
| 0.8 | 0.790 | 0.646 |
| 0.9 | 0.789 | 0.648 |
| 1.0 | 0.770 | 0.699 |

Finally, the variance-matching regularizer

$$\mathcal{L}_{\mathrm{var}}(\widetilde{Z}, \widehat{Z}) = \frac{1}{k}\sum_{a=1}^{k}\Big(\mathrm{Var}_n(\widetilde{Z}_{\cdot a}) - \mathrm{Var}_n(\widehat{Z}_{\cdot a})\Big)^2 \tag{1}$$

encourages the generated confounders to match the marginal scale of the reference confounder while preventing $\widetilde{Z}$ from collapsing to trivial or near-constant values.

We combine the regularizers into a single term,

$$\mathcal{L}_{\mathrm{reg}} \;=\; \lambda_{\mathrm{corr}} \cdot \mathcal{L}_{\mathrm{corr}} \;+\; \lambda_{\mathrm{cos}} \cdot \mathcal{L}_{\mathrm{cos}} \;+\; \lambda_{\mathrm{var}} \cdot \mathcal{L}_{\mathrm{var}},$$

where $\lambda_{\mathrm{corr}}, \lambda_{\mathrm{cos}}, \lambda_{\mathrm{var}} \geq 0$ are tuning parameters.

We include an ablation study demonstrating that regularization stabilizes training. Removing variance-matching regularization leads to less consistent recovery of latent variables across $\kappa$ levels (Table S1), whereas full regularization yields smoother behavior. Importantly, the resulting sensitivity contours and gene categorizations remain largely unchanged, indicating that regularization primarily stabilizes optimization and latent variable estimation rather than altering the final scientific conclusions.

### S6 Hyperparameters and training details

In Section S6.1, we document all the hyperparameters used in sensGAN. In Section S6.2, we describe in detail any additional training details of sensGAN not mentioned in the main text. In Section S6.4, we document the various ways we find empirically beneficial when post-processing the results after Step 2 of sensGAN (i.e., adversarially training the predictors and generator).

#### S6.1 Hyperparameters for loss functions

**Optimization and training schedule.** sensGAN alternates between refitting two predictive models and updating a generator. All components are optimized using the Adam optimizer as implemented in PyTorch, using learning rates specified below; we use `zero_grad()` only to clear accumulated gradients between backpropagation steps.

**Predictor refitting.** At each outer iteration, we refit the treatment predictor $f_D(\cdot)$ and the outcome predictor $f_Y(\cdot)$ on the current generated confounder $\widetilde{Z}$. For both predictors, `lr` denotes the optimizer learning rate, `epochs` the maximum number of training epochs, and `tol` a convergence tolerance used to stop training early when improvements fall below a threshold.

- **Treatment predictor $f_D(\cdot)$.** We train $f_D$ with the binary cross-entropy loss with logits using a learning rate `lr_cls`$= 2 \times 10^{-1}$ and at most `epochs_cls`$= 10^4$ epochs. We use a validation split `split_cls`$= 0.7$ and early stopping with patience `patience_cls`$= 200$ and tolerance `tol_cls`$= 10^{-3}$.
- **Outcome predictor $f_Y(\cdot)$.** We train $f_Y$ by maximizing the negative binomial likelihood with a learning rate `lr_nb`$= 2 \times 10^{-1}$ and `epochs_nb`$= 10^3$ epochs.

**Generator updates.** The generator $g(\cdot)$ is trained in an alternating fashion with the predictors. Training proceeds over `gen_steps`$= 10$ outer epochs. Within each outer epoch, the predictors are first refit on the current $\widetilde{Z}$, after which the generator is updated for up to `epochs`$= 100$ gradient steps using Adam with learning rate `lr_gen`$= 10^{-3}$.

**Generator loss weights.** We distinguish hyperparameters used for *predictor fitting* (above) from those used for the *generator objective*. In adversarial training, we penalize deviations from target predictive gains for $D$ and $Y$ with weights $\lambda_D = 1$ and $\lambda_Y = 1$, and apply scaling factors $s_D = s_Y = 10$ to normalize the magnitude of predictive-gain residuals. We additionally regularize the generator output to encourage structural alignment with the reference confounder via $\mathcal{L}_{\text{corr}}, \mathcal{L}_{\text{cos}}, \mathcal{L}_{\text{var}}$ with weights $\lambda_{\text{corr}} = \lambda_{\text{cos}} = \lambda_{\text{var}} = 10^{-3}$. Unless otherwise stated, these hyperparameters are fixed across experiments and chosen for optimization stability rather than performance tuning.

#### S6.2 Additional training details

**Initialization and warm start.** We initialize the generator by fitting $g(\cdot)$ to reproduce the most powerful confounder $\widehat{Z}$. We then warm-start the treatment and outcome predictors once using $\widehat{Z}$ before entering the outer training loop. No separate warmup stage is used beyond this initialization.

**Best-snapshot selection.** In practice, the alternating optimization may fail to simultaneously achieve the target predictive gains for both $D$ and $Y$ within a finite number of outer epochs. To ensure a stable and informative output in such cases, we track the generator state that attains The lowest total generator loss throughout training. Specifically, at each generator update, we record the current $\widetilde{Z}$ and associated statistics whenever the total loss improves upon the previously observed minimum. If the final training epochs do not meet the prescribed DBPR targets, we return the best-performing snapshot rather than the last iterate. This strategy prevents unstable late-stage updates from degrading the inferred confounder and provides a conservative estimate of the strongest confounder identified during training.

#### S6.3 Bilevel optimization as an alternative training strategy

The current sensGAN training procedure follows a minimax-style adversarial optimization, where the generator and predictors are updated in an alternating fashion. While this formulation is natural for adversarial learning, minimax optimization could exhibit training/numerical instability. In particular, alternating updates approximate a simultaneous game, where each component reacts only to the current state of the others, potentially leading to cyclical updates during training rather than a stable convergence.

An alternative formulation is to cast sensGAN as a bilevel optimization problem. In this setting, the predictors are treated as lower-level models that are optimized conditionally on the current latent confounder estimate, while the generator acts as the upper-level optimizer:

$$\min_G \mathcal{L}_{\text{sens}}\big(G, \theta_D^*(G), \theta_Y^*(G)\big)$$

subject to

$$\theta_D^*(G), \theta_Y^*(G) = \arg\min_{\theta_D, \theta_Y} \mathcal{L}_{\text{pred}}(G, \theta_D, \theta_Y),$$

where $\theta_D$ and $\theta_Y$ denote the treatment and outcome predictors, respectively. Conceptually, the generator no longer reacts to partially optimized predictors at each iteration. Instead, it updates while explicitly accounting for the predictors' optimal response to the current latent variable estimate. This corresponds to a hierarchical "leader-follower" formulation rather than a simultaneous adversarial game. From a coding implementation point of view, this requires sensGAN to differentiate through the lower-level optimization through unrolled optimization scheme when updating the parameters of the generator. We believe this formulation provides a more stable training strategy for sensGAN. Since the generator objective depends on predictive gain constraints, unstable predictor updates can propagate directly into the latent variable estimation procedure, leading to oscillatory behavior. Importantly however, this changes the optimization procedure rather than the scientific objective itself.

We thank the ICML reviewers for suggesting this direction. While bilevel optimization is not explored in the current manuscript, we view it as a promising extension for improving optimization stability and reproducibility in future versions of sensGAN.

##### S6.4 Post-processing of sensGAN results

**Calibration via leave-one-out predictive gains.** measured covariate. For each covariate $X_i$, we fit a leave-one-out (LOO) unadjusted NB GLM that excludes $X_i$, i.e. $(X_{-i}, D) \longrightarrow Y$, and compare it to the corresponding unadjusted model that includes all measured covariates. The normalized predictive gain for outcome $Y$, i.e. $\eta_{\mathrm{LOO}}$, is calculated based on these models, which captures the marginal predictive gain attributable to $X_i$. Repeating this procedure for all measured covariates yields an empirical distribution of $\eta$. We use this distribution as a calibration reference. Under the assumption that no latent confounder is stronger than a given measured covariate, we set the confounding strength parameter $\eta$ (i.e., the confounding-effect knob) to match or remain below $\eta_{\mathrm{LOO}}$.

**Selection for contour visualization.** For each target pair $(\eta, \kappa)$, sensGAN returns a generated confounder $\widetilde{Z}$ and the corresponding achieved predictive gains $(\eta', \kappa')$. Because alternating optimization may not exactly hit the prescribed targets, we apply a feasibility filter before constructing contour plots: we retain a run only if both gains are within a fixed tolerance,

$$|\eta' - \eta| \leq \epsilon \quad \text{and} \quad |\kappa' - \kappa| \leq \epsilon, \qquad \text{where } \epsilon = 0.1.$$

Only feasible runs are included when aggregating statistics (e.g., significance counts or p-value summaries) over the $(\eta, \kappa)$ grid; targets that fail this criterion are treated as missing and are not plotted.

**Isotonic smoothing.** To summarize trends as a function of predictive gain, we additionally apply an isotonic regression transform to obtain a monotone estimate of the relationship between $-\log_{10} p$ and the achieved treatment gain $\kappa'$. Concretely, we fit a non-increasing isotonic regression model with clipping for out-of-range inputs,

$$\widehat{m}(\cdot) \;=\; \mathrm{IsoReg}(\texttt{decreasing},\ \texttt{out\_of\_bounds} = \texttt{clip}).$$

Then, we report $\widehat{m}(\kappa')$ as the calibrated curve. This monotone post-processing reduces local noise across grid evaluations while preserving the expected directional effect.

**Refitting p-values for real data analyses.** For real-data applications (SLE in Appendix Section S10 and AD in Section 3.3), we refit the outcome models (`R:glmGamPoi`) using the inferred confounder $\widetilde{Z}$ to obtain final statistical significance measures. The refitting step yields calibrated p-values that account for overdispersion and mean-variance relationships in count data while decoupling statistical inference from the neural network training procedure.

#### S7 sensGAN pseudocode

We present the pseudocode for sensGAN in Algorithm 1 to provide a high-level overview of its steps.

**Algorithm 1** sensGAN: adversarial sensitivity analysis with predictive-gain constraints

**Input:** disease status $D \in \{0,1\}^n$, covariates $X \in \mathbb{R}^{n\times d}$, pseudobulk counts $Y \in \mathbb{N}^{n\times p}$, confounder dimension $k$, target $\eta^\star \in [0,1]$, sequence $\mathcal{K} \subset [0,1]$, significance level $\delta$, training tolerance $\epsilon$.
**Output:** worst-case confounders $\{\widetilde{Z}(\kappa,\eta)\}$, worst-case p-values $\{\widetilde{P}(\kappa,\eta)\}$, and sensitivity contours.
**Step 1A: Fit unadjusted models.**
Fit a GLM regressing $D$ on $X$ to obtain $\widehat{\pi}_{\text{un}}$.
Fit GLMs regressing $Y$ on $(X, D)$ to obtain $\widehat{\mu}_{\text{un}}$, $\widehat{\alpha}_{\text{un}}$, and $\text{SE}(\widehat{T}_{Y,\text{un}})$.
**Step 1B: Learn the most powerful confounder $\widehat{Z}$.**
Initialize treatment predictor $f_D(\cdot) : \{X, Z\} \mapsto D$.
Initialize outcome predictor $f_Y(\cdot) : \{X, D, Z\} \mapsto Y$.
Initialize generator $g(\cdot) : \{X, D, Y\} \mapsto \widehat{Z} \in (0,1)^{n\times k}$.
**repeat**
  Train $f_D(\cdot)$ with current $\widehat{Z}$ to minimize $\mathcal{L}_{\text{BCE}}$.
  Train $f_Y(\cdot)$ with current $\widehat{Z}$ to minimize $\mathcal{L}_{\text{NB}}$, using $\widehat{\alpha}_{\text{un}}$.
  Train $g(\cdot)$ to minimize $\mathcal{L}_L = \lambda_{\text{BCE}}\mathcal{L}_{\text{BCE}} + \lambda_{\text{NB}}\mathcal{L}_{\text{NB}}$.
**until** convergence
Compute maximal DBPRs $(\widehat{R}^2_D, \widehat{R}^2_Y)$ using $\widehat{Z}$.
**Step 2: Learn worst-case confounders under predictive-gain constraints.**
**for** fixed $\eta^\star$ and each $\kappa \in \mathcal{K}$ **do**
  **repeat**
    Generate current confounder $\widetilde{Z} = g(X, D, Y)$.
    Compute normalized predictive gains $\kappa' = \widetilde{R}^2_D / \widehat{R}^2_D$ and $\eta' = \widetilde{R}^2_Y / \widehat{R}^2_Y$.
    Train $f_D(\cdot)$ with current $\widetilde{Z}$ to minimize $\mathcal{L}_{\text{BCE}}$.
    Train $f_Y(\cdot)$ with current $\widetilde{Z}$ to minimize $\mathcal{L}_{\text{NB}}$, using $\widehat{\alpha}_{\text{un}}$.
    Train $g(\cdot)$ to minimize $\mathcal{L}_S + \mathcal{L}_P + \mathcal{L}_{\text{reg}}$ using $\delta$, $\text{SE}(\widehat{T}_{Y,\text{un}})$, $(\widehat{R}^2_D, \widehat{R}^2_Y)$, and targets $(\kappa, \eta^\star)$.
  **until** $|\kappa' - \kappa| \le \epsilon$ and $|\eta' - \eta^\star| \le \epsilon$
  Save $\widetilde{Z}$ and the corresponding worst-case p-values for all $p$ genes.
**end for**

### S8 Data simulation details

In Sections S8.1 and S8.2, we describe how we simulate a binary and continuous-valued latent confounder $Z$, respectively. In Section S8.3, we document our crawler, which is used to compute the reference sensitivity contours. In Section S8.4, we document the results of how our crawler performs in our simulation setting.

#### S8.1 Binary $Z$ simulation

We first describe the simulation used to generate the results shown in Figures 4 and 4 and Table 1 in the main text, where we demonstrate that sensGAN is able to accurately estimate the latent confounders based on $(\kappa^*, \eta^*)$ and that the genes are appropriately partitioned into different categories based on whether or not they are associated with the latent confounders. Here, we simulate the measured covariate matrix $X \in \mathbb{R}^{n\times d}$ and the latent binary confounder $Z \in \mathbb{R}^{n\times k}$ independently. Then, we simulate the treatment $D \in \mathbb{R}^{n\times 1}$ from a Bernoulli distribution with a logit link, while controlling for the variance contributed by terms involving $X$ and $Z$. With the Negative Binomial GLM link function, we simulate the mean count matrix of scRNA-seq pseudocounts $Y \in \mathbb{R}^{n\times p}$. Then, we simulate the overdispersion rate to be a normal distribution centered around a constant vector $C \in \mathbb{R}^{p\times 1}$. Our simulated data can be compactly denoted as,

$$\begin{aligned} &D \sim \text{Bernoulli}(\sigma(XB_D + Z\Gamma_D)) \\ &Y \sim \text{NB}(\log(\mu), \alpha), \text{ where } \log(\mu) = DT_Y + XB_Y + Z\Gamma_Y, \end{aligned}$$

where $\sigma(\cdot)$ denotes the sigmoid function.

With different combinations of $T_Y \in \mathbb{R}^{1\times p}$ and $\Gamma_Y \in \mathbb{R}^{k\times p}$, we simulate 3 categories of genes with varied significance profiles:

- **Category i ("Significant")**: Genes significantly associated with $D$ but not with $Z$. Their expression remains significantly associated with $D$ both with and without adjusting for $Z$. They are later referred to as the (always) significant genes.
- **Category ii ("Nullified")**: Genes significantly associated with $Z$ but not with $D$. These genes appear significant without adjusting for $Z$ but become insignificant once $Z$ is included in the model. They are later referred to as the nullified genes.
- **Category iii ("Insignificant")**: Genes not significantly associated with either $D$ or $Z$. They remain insignificant with and without adjustment for $Z$. They are later referred to as the (always) insignificant genes.

In this simulation, we set $n = 100$, $p = 100$, $d = 4$, $k = 1$, $C = 0.5$.

#### S8.2 Continuous $Z$ simulation for contour validation

Next, we describe the simulation used to generate Figure 5, which validates the contours of each gene. This simulation experiment validates the contour curves generated by sensGAN. Most simulation settings are similar to the above. The one difference is that we now simulate the continuous latent confounder $Z \in (0,1)^{n\times k}$, where each entry is drawn independently and identically distributed from a uniform distribution. As mentioned in the main text, we use a continuous $Z$ in this simulation to have finer control over predictive gains. In this experiment, we only simulate genes in **Category ii**, set $n = 100$, $p = 10$, $d = 4$, $k = 1$, $C = 0.5$. We use $p = 10$ so our contour validation works better empirically (see Appendix Section S8.3 for details of our crawler) and so we can restrict attention to genes that are associated with $Z$ but not with $D$. Although Category ii genes are simulated to be nullified by adjustment for $Z$, in practice, not all such genes are empirically nullified, since the outcome predictive gain of $Z$ is constrained by the target $\eta$ and finite-sample randomness can obscure marginal significance changes.

#### S8.3 Crawler-based confounder generation.

To validate the contour generated by sensGAN, we implement a crawler-style worst-case search that directly explores the space of continuous confounders $Z \in [0,1]^n$. This procedure is used to generate the *reference contours* shown in Figure 6 of the main text.

The crawler iteratively proposes candidate confounders using a diverse set of heuristics designed to cover a broad range of plausible confounding structures:

- **Independent random draws**: Sampling $Z$ i.i.d. from Uniform$(0,1)$.
- **Perturbations of the true confounder**: Adding noise to the simulated ground-truth $Z$ to explore nearby confounding directions.
- **Rank-based transformations of $D$**: Constructing $Z$ from the ranks of the treatment assignment, optionally with added noise.
- **Rank-based transformations of $Y$**: Using high-variance genes in $Y$ to induce confounders correlated with outcome structure.
- **Principal components of $(Y, D)$**: Extracting leading principal components from the concatenated matrix of outcomes and treatment.
- **Jittered refinements of previous candidates**: Locally perturbing previously evaluated confounders to refine coverage of promising regions.

Each proposed confounder $Z$ is evaluated by refitting adjusted models for $D$ and $Y$ and computing the corresponding deviance-based partial $R^2$ values. These are normalized to obtain predictive-gain coordinates $(\kappa, \eta)$. Candidates are then archived on a predefined $(\kappa, \eta)$ grid, and for each grid cell, the crawler retains the confounder that minimizes the number of significant genes, yielding an explicit approximation to the worst-case confounding scenario.

##### S8.4 Contour construction with crawler-based confounder.

To construct the reference contours, we discretize both $\kappa$ and $\eta$ into 11 equally spaced bins over $[0, 1]$, resulting in an $11 \times 11$ grid (Fig. S1). Only confounders whose achieved predictive gains fall within the corresponding bin are used to summarize that grid cell.

Table S2 reports the number of candidate confounders evaluated by the crawler that fall into each $\kappa$ bin ($\pm 0.05$), illustrating coverage of the predictive-gain space.

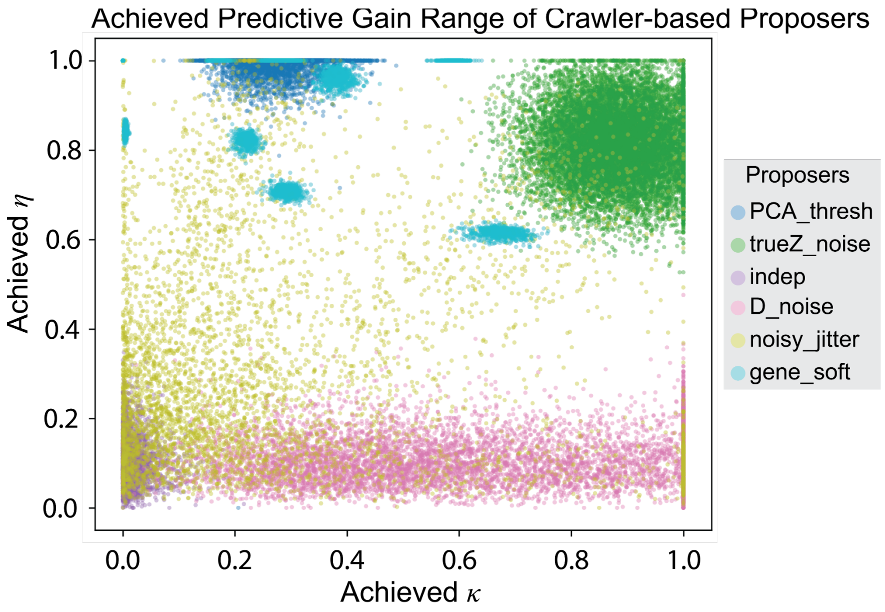


Figure S1: **Coverage of the $(\kappa, \eta)$ predictive-gain space achieved by the crawler-based proposers.** Each point corresponds to a candidate confounder $Z$ proposed by the crawler and evaluated by its normalized predictive gains for $D$ ($\kappa$) and $Y$ ($\eta$). Different proposal heuristics populate distinct regions of the grid, illustrating the heterogeneous and uneven coverage induced by direct search. This visualization motivates the need for sensGAN to more systematically and continuously explore the predictive-gain space.

Table S2: Number of crawler-generated confounders in each $\kappa$ bin ($\pm 0.05$).

| $\kappa$ bin | 0.0 | 0.1 | 0.2 | 0.3 | 0.4 | 0.5 | 0.6 | 0.7 | 0.8 | 0.9 | 1.0 |
|---|---|---|---|---|---|---|---|---|---|---|---|
| Count | 42 | 72 | 60 | 37 | 21 | 30 | 32 | 27 | 20 | 5 | 8 |

#### S9 Additional results: Simulation

**Gene-level nullification contingency analysis.** To evaluate sensGAN at the gene level by comparing per-gene sensitivity contour curves, we further summarize gene-level behavior with a contingency table that classifies genes as always significant, nullified, or always insignificant (Table S3). The contingency analysis shows that sensGAN accurately recovers the nullification status for the majority of genes, with consistent agreement across all three categories.

**Effect size versus correlation structure.** To further investigate the interpretation of sensitivity, we perform an additional simulation study examining the interplay between effect size and correlation structure. Building upon the simulation framework in Section 3.1, the genes here associated with the latent confounder (but not with the disease variable) are divided into two subgroups: (i) correlated genes with larger confounder effect sizes (mean $\approx 3$, std $\approx 2$), and (ii) independent genes with smaller confounder effect sizes (mean $\approx 1.5$, std $\approx 2$). We then compare their sensitivity contour behaviors under sensGAN. We observe that

Table S3: **Gene-level nullification agreement between the crawler reference and sensGAN.** The table reports the number of genes assigned to each significance category by sensGAN, stratified by the corresponding categories identified by the crawler-based reference. sensGAN correctly identifies 3 out of 4 genes that are nullified under worst-case confounding in the crawler analysis.

| Method | Category | Significant | Nullified | Insignificant |
|---|---|---|---|---|
| sensGAN | Significant | 1 | 0 | 0 |
| | Nullified | 0 | 3 | 1 |
| | Insignificant | 0 | 0 | 5 |

sensGAN preferentially nullifies correlated features, even when effect sizes are comparable. In particular, genes in subgroup (i) exhibit earlier sensitivity contour crossing points, becoming non-significant under smaller predictive gain requirements. These results suggest that sensGAN captures not only marginal effect size, but also the extent to which signals can be jointly explained by a shared latent structure. This behavior is consistent with many biological settings, where genes operate through coordinated pathways and shared regulatory programs.

### S10 Additional results: Systemic lupus erythematosus analysis

Systemic lupus erythematosus (SLE) is an autoimmune disease predominantly affecting women and individuals of Asian, African, and Hispanic descent (Perez et al., 2022). Here, the authors use multiplexed single-cell RNA sequencing to capture the complexity of immune cell populations and systematically profile the composition and transcriptional states of immune cells in a large multiethnic cohort. Our goal for this analysis is to demonstrate the rigor of sensGAN: withholding key donor covariates yields results similar to those from a differential expression analysis that uses them.

To remove genes with small variations, we preprocess the single-cell data by selecting the top 2,000 highly variable genes (HVGs) within each cell type, computing pseudobulk gene expression vector for each donor, and normalizing counts to the total library size per cell. With a focus on the T4 cell type, we aggregate single-cell expression profiles by summing counts across cells from the same subject, yielding a gene-level pseudobulk count matrix. Then, we remove genes with over 90% zero counts across subjects, retaining those expressed in at least 10% of donors for downstream analysis. To ensure method stability, we also remove genes whose counts do not converge in the naive GLM Negative Binomial Model. In total, the dataset includes $n = 256$ subjects (158 cases and 98 controls). For each subject, the case–control variable represents systemic lupus erythematosus (SLE) status, and the final number of retained genes is $p = 475$.

In the proof-of-principle analysis, we only provide the method with sex ($d = 1$) while holding out other measured covariates, including ethnicity and the sequencing batch information. To make sure the method captures biologically meaningful confounders, we decide to search for confounders ($k = 3$) with a wider range of confounding effects by letting $\kappa$ and $\eta$ be equal to each other and vary together, i.e., letting $Z$'s predictive gains on $Y$ be equal to $Z$'s predictive gains on $X$. With a series of $(\kappa, \eta)$ values from 0.2 to 1, with a step size of 0.05, the method identifies a series of confounders. We then perform adjusted differential expression analyses with confounders of varying strengths.

First, we analyze the significance profiles of genes under varying levels of confounder strengths. In the sensitivity contour plot (Fig. S2A), mean contour curves and 95% confidence intervals are colored by genes that are always significant, nullified, and always insignificant. Then, we calculate the exact $\kappa$ at which the gene become insignificant by extrapolating on the stepwise contour curve (Fig. S2B). Shown by the bar graph of the number of differentially expressed genes in the $\kappa$s, we also observe that larger $\kappa$ generally corresponds to fewer differentially expressed genes (Fig. S2C). These results show that the method can identify a series of confounders with varying strengths, and that more significant DEGs in the original model become insignificant as $\kappa$ increases.

Then, we analyze the DEG sets across 3 methods to understand how sensGAN compares to existing methods while applying to real data. We use the Gamma-Poisson regression model to identify the DEGs associated with SLE, adjusting for different sets of covariates. First, for the sensGAN method, we adjust for sex and the sensGAN-learned confounders. Second, for the all-covariate method, we adjust for all measured covariates. Third, for the unadjusted method, we do not adjust for latent confounding but only adjust for

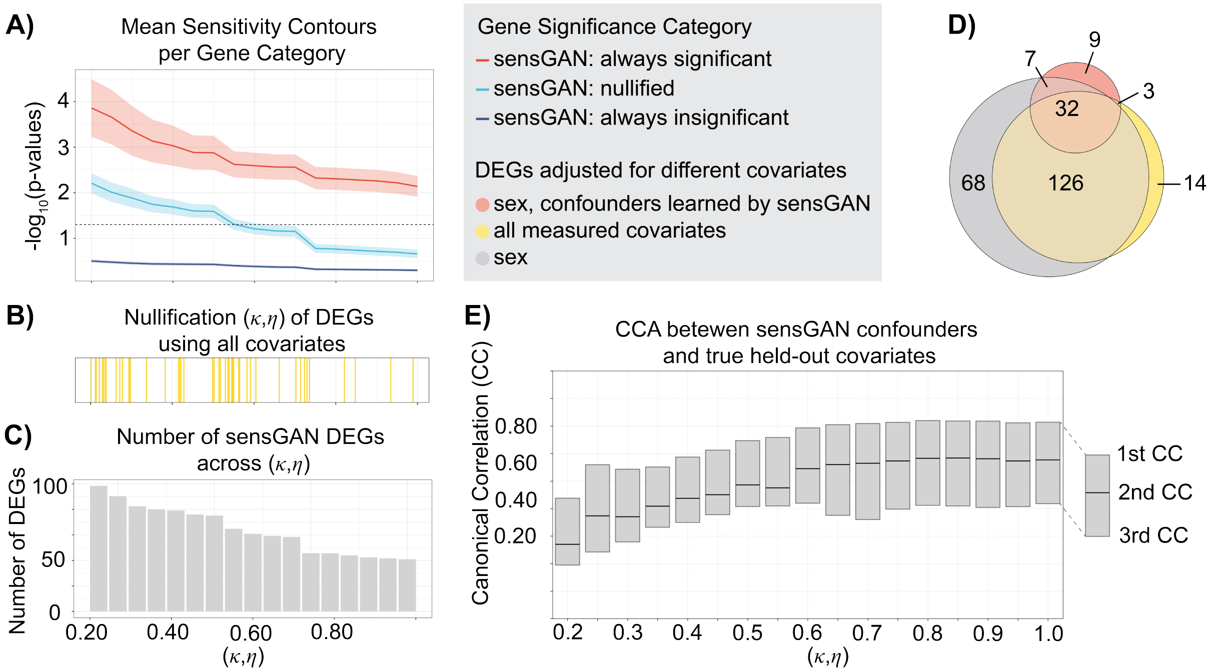


Figure S2: **sensGAN identifies robust SLE DEGs under increasing latent confounding and learns biologically meaningful confounders aligned with held-out covariates.** **A)** The sensitivity diagnostic contour plot with ($\kappa$, $\eta$), where $\kappa = \eta$, varying from 0.2 to 1. **B)** The $\kappa$ and $\eta$ values where genes are nullified, extrapolated from the stepwise curve. **C)** The barplot of the number of DEGs at varying ($\kappa$, $\eta$) pairs. **D)** The Venn plot of significant DEGs calculated by the GLM Gamma-Poisson model adjusting for different covariates. **E)** The canonical correlations between the sensGAN confounders at varying levels of strength and the held-out covariates.

sex. We present the results as a Venn diagram, in which circle areas correspond to the total number of DEGs identified while adjusting for different sets of covariates (Fig. S2D). The unadjusted DEG set is the largest, partially because it only adjusts for sex. The all-covariate DEG set is smaller than the unadjusted DEG set and larger than the sensGAN DEG set, partially due to its adjustment of other measured covariates. The sensGAN DEG set only includes the genes that are always significant with varying levels of confounding strengths; this is the smallest set and most of its DEGs are also identified by the other methods.

Lastly, we summarize the relationships between the confounders ($k = 3$) learned by sensGAN and the held-out covariates ($d' = 4$) using canonical correlation analyses (CCA) (Fig. S2E). Every bar corresponds to 3 canonical correlation (CC) values while comparing the held-out covariates and the learned confounders. We observe that all CCs increase as the confounding strengths increase from 0.2 to 1. Based on this observation, we believe that the method can learn a series of biologically meaningful confounders that become increasingly similar to the held-out covariates as we allow stronger confounding effects.
